# Dengue and Zika virus infection are impaired by small molecule ER proteostasis regulator 147 in an ATF6-independent manner

**DOI:** 10.1101/2020.05.15.098624

**Authors:** Katherine M. Almasy, Jonathan P. Davies, Samantha M. Lisy, Reyhaneh Tirgar, Sirena C. Tran, Lars Plate

**Author notes:** Corresponding author: Lars Plate, Departments of Chemistry and Biological Sciences, Vanderbilt University, Nashville, TN 37340.

## Abstract

Flaviviruses, including Dengue and Zika, are widespread human pathogens, however, no broadly active therapeutics exist to fight infection. Here, we establish the recently discovered pharmacologic modulator of ER proteostasis **147** as an effective host-centered antiviral strategy. Compound **147** reduces infection by attenuating viral replication without causing toxicity in host cells. **147** is a preferential activator of the ATF6 pathway of the unfolded protein response, which requires targeting of cysteine residues primarily on protein disulfide isomerases (PDIs). We find that the antiviral activity of **147** is independent of ATF6 induction but does require modification of reactive thiols on protein targets. Targeting PDIs using RNAi and other PDI small molecule inhibitors was unable to recapitulate the antiviral effects, suggesting additional identified protein targets of **147** may mediate the activity. Importantly, **147** can impair infection of multiple strains of Dengue and Zika virus, indicating that it is suitable as a broad-spectrum antiviral agent.

## INTRODUCTION

Flaviviruses are significant human pathogens that cause widespread mortality around the world. The genus encompasses several arthropod-borne viruses including Dengue (DENV), Yellow Fever (YFV), and Zika (ZIKV) (Bhatt et al., 2013; Daep et al., 2014; Weaver et al., 2016). These viruses pose serious public health threats, particularly as increased global travel and climate change have caused an increase in the spread of both the virus and their mosquito vectors (*A. albopictus* and *A. aegypti*) (Tatem et al., 2012; Whitehorn and Yacoub, 2019). It is estimated that 53% of the world population lives in areas suitable for Dengue transmission resulting in 50-100 million Dengue infections per year, of which 500,000 proceed to Dengue hemorrhagic fever and 22,000 are fatal (Bhatt et al., 2013; Messina et al., 2019). ZIKV infection has been linked to serious neurological defects, including Guillain-Barré syndrome and microcephaly in newborns (Mendez et al., 2017). While ZIKV diagnoses were relatively rare until the early 2000s, the 2015–2016 pandemic in the Americas highlighted the potential for rapid spread of the disease vectors (Weaver et al., 2016).

Vaccines currently exist for only a limited range of flaviviruses (Collins and Metz, 2017). The first DENV vaccine, Dengvaxia, was approved by the FDA in early 2019 (Thomas and Yoon, 2019). However, its use remains limited to children with confirmed prior infection. In addition, no post-exposure therapeutic options are available for patients infected with these viruses; current treatments only attempt to alleviate the symptoms (Bernatchez et al., 2020; Lim, 2019; Lim et al., 2013). Antiviral strategies often focus on directly targeting viral proteins. While molecules inhibiting for instance the flavivirus RNA-dependent RNA polymerase NS5 or the protease NS3 have been identified, the high mutation rate of the virus allows for resistance to be developed quickly (Bernatchez et al., 2020; Carrasco-Hernandez et al., 2017; Garcia et al., 2017). One increasing area of exploration for alternative therapeutic approaches is the targeting of host factors that are critical for virus propagation (Kaufmann et al., 2018). As these proteins are not under genetic control of the virus, development of resistance is much less likely (De Clercq, 2007; Geller et al., 2013, 2007). In addition, host-targeted therapeutics should be effective as broad-spectrum antivirals instead of targeting a single virus (Aviner and Frydman, 2020; Taguwa et al., 2019, 2015).

Flaviviruses contain a genome of positive-sense, single stranded RNA (+ssRNA) approximately 11kb in size. The viruses replicate and assemble around the endoplasmic reticulum (ER) membrane (Apte-Sengupta et al., 2014). As the genome is translated after entry into the cell, the single polyprotein is inserted into the ER membrane and is co- and post-translationally processed into three structural (capsid, pre-membrane, and envelope) and seven nonstructural (NS1, NS2A, NS2B, NS3, NS4A, NS4B, NS5) proteins (Perera and Kuhn, 2008). The host proteostasis network has been implicated in the maintenance of individual viral proteins and virus evolution (Aviner and Frydman, 2020; Phillips et al., 2017; Ravindran, 2018; Reid et al., 2018). The proteostasis network is comprised of chaperones, co-chaperones and other protein quality control factors that control protein folding, assembly, post-translational modification, trafficking, and degradation. Genetic screens have identified many proteostasis factors as essential for DENV replication, including several components of the oligosaccharyl transferase (OST) complex, as well as the ER membrane protein complex (Lin et al., 2019; Marceau et al., 2016; Ngo et al., 2019; Savidis et al., 2016). Aside from promoting biogenesis of viral proteins, these components also have additional roles in organizing viral replication centers at the ER membrane (Marceau et al., 2016; Ngo et al., 2019; Rothan and Kumar, 2019). Additionally, diverse cellular chaperone and co-chaperone systems are required for all stages of the viral life cycle, including virus entry and disassembly, folding of individual viral proteins, as well as assembly and egress of new virions (Fischl and Bartenschlager, 2011; Heaton et al., 2016; Taguwa et al., 2019). Pharmacologic inhibition of proteostasis factors, including the OST as well as cytosolic Hsp70 and Hsp90 chaperones, has been shown to be an effective strategy to reduce flavivirus infection in cell models (Heaton et al., 2016; Howe et al., 2016; Puschnik et al., 2017; Taguwa et al., 2019, 2015; Yang et al., 2020).

As flavivirus replication and polyprotein translation occurs at the ER membrane, infection results in expansion of the ER and remodeling of the ER proteostasis environment (Ravindran et al., 2016; Rothan and Kumar, 2019; Stohlman et al., 1975). DENV infection leads to modulation of the unfolded protein response (UPR), the adaptive stress-response that remodels the ER proteostasis environment to counter stress caused by accumulation of misfolded proteins (Peña and Harris, 2011; Perera et al., 2017). The UPR consists of three overlapping but distinct signaling branches downstream of the stress-sensing receptors IRE1 (inositol-requiring enzyme 1), ATF6 (activating transcription factor 6), and PERK (Protein kinase R-like ER kinase) (Shoulders et al., 2013; Walter and Ron, 2011). The first two branches primarily control the upregulation of chaperones and other proteostasis factors to expand the protein folding capacity of the ER. The PERK branch is responsible for translational attenuation via phosphorylation of eukaryotic initiation factor 2 alpha (eIF2α) and subsequent activation of the integrated stress response. DENV has been shown to upregulate the IRE1/XBP1s and ATF6 branches of the UPR, while suppressing activation of the PERK branch (Peña and Harris, 2011). The importance of UPR-dependent ER proteostasis remodeling for flavivirus infection has only been explored in knockout cell models. For example, the virus was less capable of propagating in IRE1 -/- MEF cells. In contrast, PERK knockout resulted in increased viral titers, while ATF6 knockout did not impact viral propagation. (Peña and Harris, 2011).

Given the dependencies of DENV and other flaviviruses on UPR modulation and an enhanced ER proteostasis environment in the host cells during infection, we sought to explore whether these dependencies can be perturbed to impair viral infection. Pharmacologic remodeling of ER proteostasis pathways has become an attractive strategy at correcting imbalances associated with diverse phenotypes related to protein stress and misfolding without affecting endogenous protein maturation or causing toxicity (Blackwood et al., 2019; Chen et al., 2014; Cooley et al., 2014; Kroeger et al., 2018; Plate et al., 2016). In particular, compound **147**, which was developed as a preferential activator of the ATF6 pathway, has broadly beneficial effects at reducing secretion of amyloidogenic proteins and protecting against oxidative organ damage from ischemia/reperfusion (Blackwood et al., 2019; Plate et al., 2016).

Here, we demonstrate that ER proteostasis remodeling by compound **147** serves as an effective strategy to reduce flavivirus infection. We determine that the **147** mediated reduction in DENV replication and assembly is surprisingly not mediated through activation of ATF6. Instead, the activity is mediated by upstream covalent modifications of protein targets by **147**. Prior work identified protein disulfide isomerases (PDIs) as critical targets for the **147**-dependent ATF6 activation (Paxman et al., 2018), however, our data suggests PDIs are not responsible for the antiviral activity. Finally, we show that **147** treatment can reduce proliferation of multiple DENV serotypes and several ZIKV strains, demonstrating that the pharmacologic agent could be a broadly effective strategy against flaviviruses and other viruses that depend on ER proteostasis processes.

## RESULTS

### A Selective Modulator of the ATF6 Pathway Impairs Dengue Virus Infection

DENV is known to activate the UPR in infected host cells and was previously found to specifically upregulate the ATF6 and IRE1/XBP1s signaling arms, which transcriptionally upregulate ER protein quality control factors and adjust proteostasis capacity (**Fig. 1A**) (Peña and Harris, 2011; Shoulders et al., 2013). To confirm the activation of ER proteostasis pathways by DENV, we performed an infection time-course with DENV serotype 2 (DENV2) in Huh7 liver carcinoma cells and measured transcript levels of an ATF6-regulated gene (*PDIA4*) and an XBP1s-regulated gene (*ERDJ4*) by quantitative RT-PCR. Both *PDIA4* and *ERDJ4* transcripts exhibited a time-dependent induction in response to DENV2 infection (**Fig. 1B**). A smaller but similar time-dependent upregulation of ATF6 and XBP1s-regulated protein products could be observed by western blot (**Fig. S1A**). To corroborate these findings, we performed quantitative proteomics analysis of Huh7 cells at 24 and 36 hours post infection (hpi) with DENV2 to quantify global changes in protein expression (**Fig. S1B-C, Table S1**). We found that DENV2 activated specific factors involved in ER proteostasis maintenance, such as DNAJC10, BiP, and HYOU1. We also filtered previously defined genesets consisting of ∼20 specific genes for each of the UPR signaling branches to quantify the cumulative activation of ER proteostasis pathways (**Fig. 1C**) (Grandjean et al., 2019). This analysis indicated a mild stimulation of ATF6 and XBP1s-regulated proteins, while PERK-regulated proteins remained largely unchanged. In contrast, we did not observe upregulation of cytosolic proteostasis factors that are under control of the heat shock response (HSR) (Vabulas et al., 2010). Overall, our results confirm that DENV2 preferentially remodels ER proteostasis pathways through ATF6 and IRE1/XBP1s-dependent upregulation of specific chaperones and protein folding factors.

**Figure 1 – (with 1 supplement).**
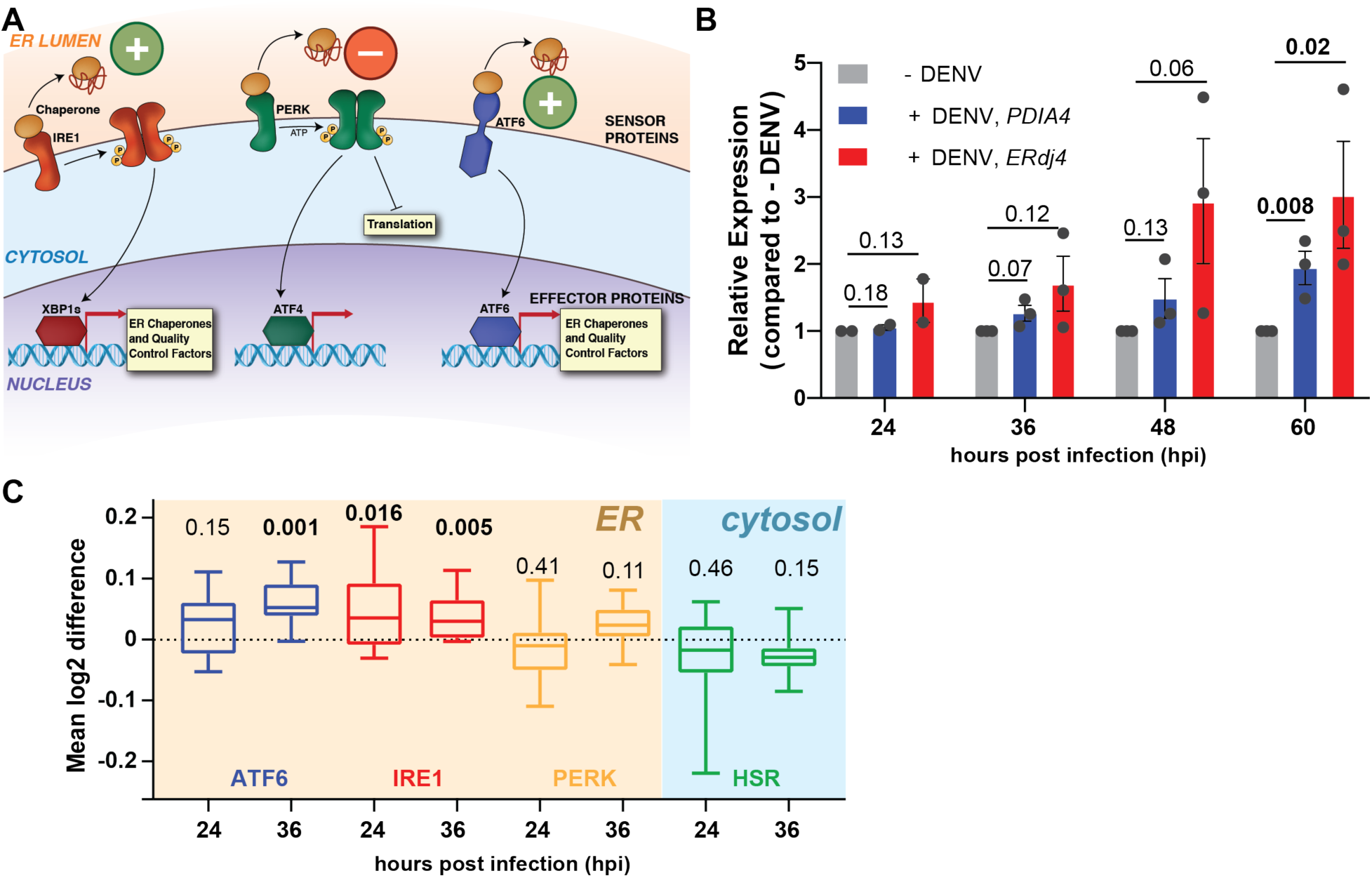
DENV infections activates the ER Unfolded Protein Response (UPR). A. Schematic overview of the UPR showing the three signaling branches: IRE1/XBP1s, PERK, and ATF6. Prior work showed that DENV increased activity of the IRE1/Xbp1s and ATF6 branches, while simultaneously preventing activation of the PERK branch (Pena and Harris, 2011). B. Time course of qPCR data showing transcriptional upregulation of an IRE1/XBP1s target (*ERdj4*) and an ATF6 target (*PDIA4*) in response to DENV activation (MOI 3) in Huh7 cells. Error bars shows SEM from 2 to 3 biological replicates and p-values from two-tailed unpaired student t-tests are displayed. C. Box plots of proteomics data showing the aggregate upregulation of IRE1/XBP1s and ATF6 protein targets at 24 and 36 hpi. PERK and cytosolic heat-shock response (HSR) targets are not affected. Huh7 cells were infected with DENV (MOI 3) for 24h or 36h. Genesets for UPR and HSR pathways were defined based on prior transcriptional profiles (Grandjean et al., 2019). p-values from two-tailed Wilcoxon signed rank tests are indicated. Cell-wide proteomics data comparing DENV infected to non-infected Huh7 cells is shown in **Fig. S1B-C**.

Considering the dependencies of DENV and other flaviviruses on UPR activation, we determined the impact of the proteostasis regulators **147** (a preferential activator of the ATF6 branch), and Ceapin-A7 (**Cp7**) (a selective inhibitor of the ATF6 branch) on DENV infection (**Fig. 2A**). We pre-treated Huh7 cells with the respective compounds 16h prior to infection by DENV2 to induce or inhibit UPR-dependent proteostasis remodeling, and we re-treated immediately after infection. Quantification of DENV viral titers by a focus forming assay demonstrated a significant reduction in infection with compound **147** at 24 and 36 hpi (**Fig. 2B**). In contrast, **Cp7** treatment only resulted in a small reduction in viral titers at 24 hpi but the infection recovered at 36 hpi. Cell viability experiments indicated that the compounds did not induce a large amount of cell toxicity in Huh7 cells consistent with previous results in other cell lines (**Fig. S2A**) (Plate et al., 2016). We also conducted dose-response studies with **147** demonstrating that the compound was effective at reducing DENV titers at an IC50 of approximately 1 µM (**Fig. S2B**). We observed that cell viability was maintained above 90% at this concentration (**Fig. S2C**). These results demonstrate that modulation of the ER proteostasis network with the preferential ATF6 activator **147** could represent an effective strategy to impair DENV2 infection.

**Figure 2 – (with 1 supplement).**
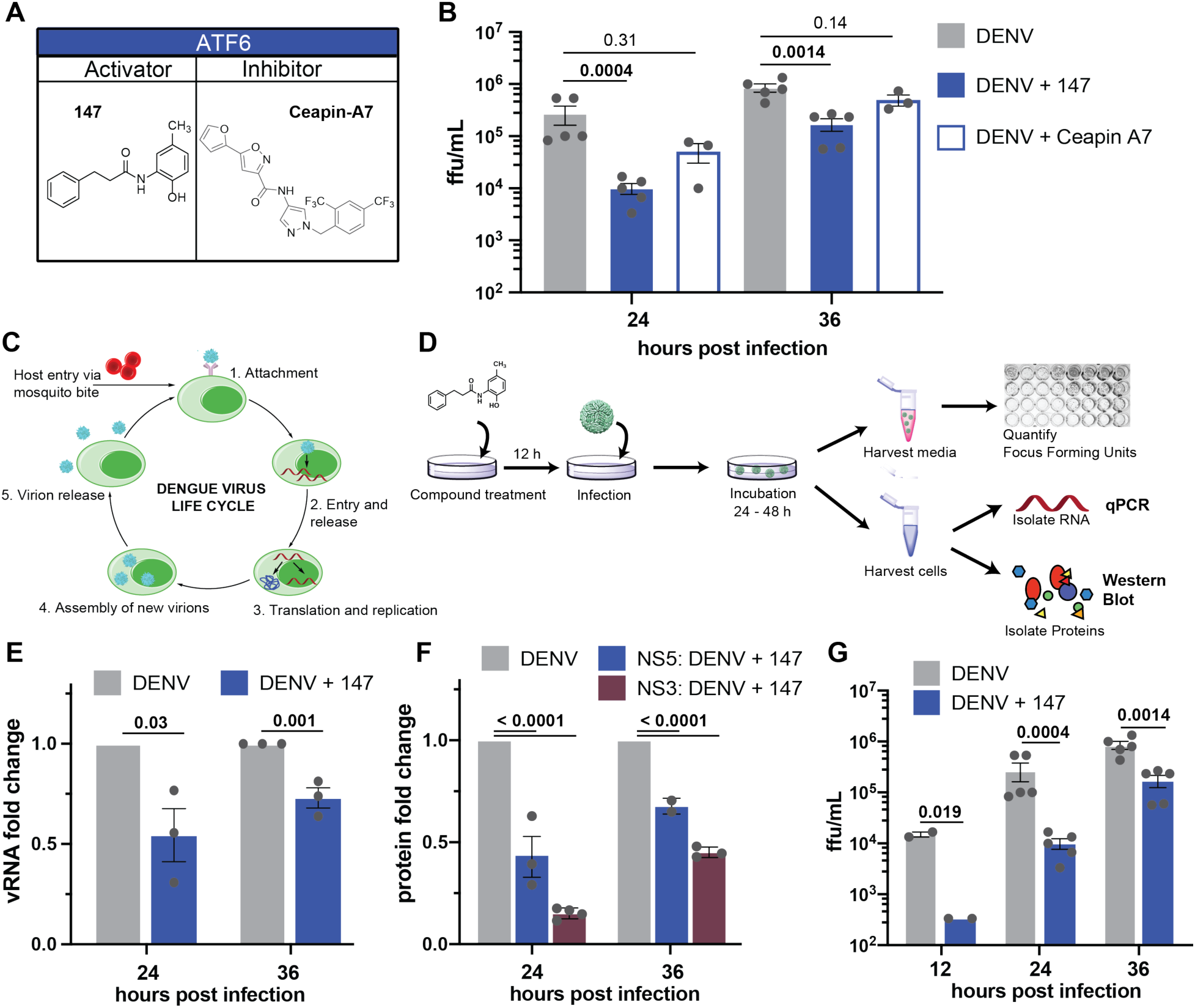
Treatment with small molecule 147 reduces viral RNA, protein, and titer levels. A. Chemical structures of small molecule ATF6 activator **147** and ATF6 inhibitor **Ceapin-A7 (Cp7)**. B. Focus forming assay to quantify changes in Dengue infection in response to treatment with ATF6 modulator compounds. Huh7 cells were treated with compounds, and 16 h later infected with DENV-2 BID-V533 at a MOI of 3 for 3 hours. Media and treatments were replaced, and cells and media were harvested at indicated timepoints and virus focus forming units (ffu) quantified. Errors bars show SEM and p-values from ratio paired t-tests are indicated. C. Schematic of the flavivirus life cycle. After attachment of the virus to a cellular receptor and clathrin-mediated endocytosis, fusion of the envelope protein with the endosome causes release of the capsid and RNA genome. The positive sense genome is then translated, providing the machinery to begin forming replication pockets at the ER membrane. Once sufficient proteins are available, new virions are assembled, trafficked through the Golgi network, and released into the extracellular space. D. Experimental workflow for quantifying viral RNA, viral proteins and titers. After pretreatment with compound **147**, cells are infected and left for harvesting at later timepoints. Media samples are taken to quantify ffu; cellular samples taken to quantify vRNA by qPCR or viral proteins by Western Blot/proteomics analysis. E. Bar graph showing reduction in viral RNA in response to **147** treatment as outlined in **D** 24 and 36 hpi. Error bars correspond to SEM of 3 biological replicates and p-values from unpaired t-tests are shown. F. Bar graph showing reduction in NS3 and NS5 viral protein levels in response to **147** treatment as outlined in **D** 24 and 36 hpi. Error bars correspond to SEM from 2 to 3 biological replicates and p-values from unpaired t-tests are shown. G. Bar graph showing reduction DENV viral titers in response to **147** treatment as outlined in **D** 24 and 36 hpi. Error bars correspond to SEM from 2 to 5 biological replicates and p-values from ratio paired t-tests are shown.

The ER plays critical roles in several stages of the viral life cycle including replication of viral RNA at replication centers on the cytosolic side of the ER membrane, translation and proteolytic processing of the viral polyprotein in the ER membrane, as well as the folding, assembly and secretion of new virions (Li et al., 2015; Neufeldt et al., 2018). To determine at what stage in the viral life cycle the compound treatment impaired viral propagation, we investigated the impact of compound **147** on viral RNA (vRNA) and protein levels (**Fig. 2C-D**). Quantification of vRNA by qPCR indicated a modest but significant reduction of 50% at 24 hpi, which was attenuated to 30% reduction at 36 hpi (**Fig. 2E**). Western blot quantification of NS3 and NS5 viral protein in Huh7 cell lysates from DENV2 infected cells showed a 35 - 80 % reduction in viral non-structural proteins in response to **147** treatment (**Fig. 2F, Fig. S2D**). This highlighted that the ER proteostasis regulator exerted a more pronounced effect on viral protein production relative to replication of vRNA. In contrast, infectious viral titers showed a far greater reduction reaching 98% at 12 hpi and sustaining 80% reduction at 36 hpi (**Fig. 2G**). The intensified effect of **147** on viral titers compared to vRNA and protein levels suggests that the compound predominantly acts on a stage subsequent to translation of viral proteins, which would be consistent with disruption of viral maturation or secretion pathways through modulation of ER proteostasis. Lastly, we confirmed that **147** did not impair viral entry. Omission of the pretreatment prior to infection and treatment of Huh7 cells with the compound only after infection still resulted in a comparable reduction in viral protein and titers, confirming that the compound must act at a post-entry stage (**Fig. S2E-G**).

### Inhibition of DENV infection by 147 is only partially dependent on ATF6

Given that flavivirus infection activates the ATF6 pathway, it seemed surprising that pre-activation could reduce viral propagation. We therefore sought to investigate whether ATF6 activation and induction of ATF6-targeted proteostasis factors was required for the **147**-dependent inhibition of DENV infection. We first confirmed by qPCR and quantitative Western blot analysis that treatment with **147** activated the ATF6 regulated gene *HSPA5* (*BiP*) in Huh7 cells (**Fig. S3A**). To probe the impact of ATF6 activation on the compound activity, we took advantage of the ATF6 inhibitor **Cp7**, which inhibits ATF6 (Gallagher et al., 2016; Gallagher and Walter, 2016). **Cp7** mediates a neomorphic interaction between ATF6 and the peroxisomal membrane protein ABCD3, keeping ATF6 in a trafficking-incompetent oligomeric state, which cannot be activated by **147** (**Fig. 3A**) (Torres et al., 2019). We confirmed by qPCR and Western blot that co-treatment with **147** and **Cp7** could fully attenuate the **147**-dependent induction of ATF6 target genes in Huh7 cells (**Fig. S3A**). We then investigated the addition of **Cp7** on the **147**-mediated reduction of DENV propagation. Co-treatment of **Cp7** did not diminish the reduction in vRNA or NS3 protein (**Fig. 3B-C**). Furthermore, the addition of **Cp7** only partially recovered the DENV2 viral titers at 24hpi (**Fig. 3D**). These results highlight that the reduction in viral titers could not be fully attributed to the **147**-mediated induction of ATF6 target genes.

**Figure 3 – (with 1 supplement).**
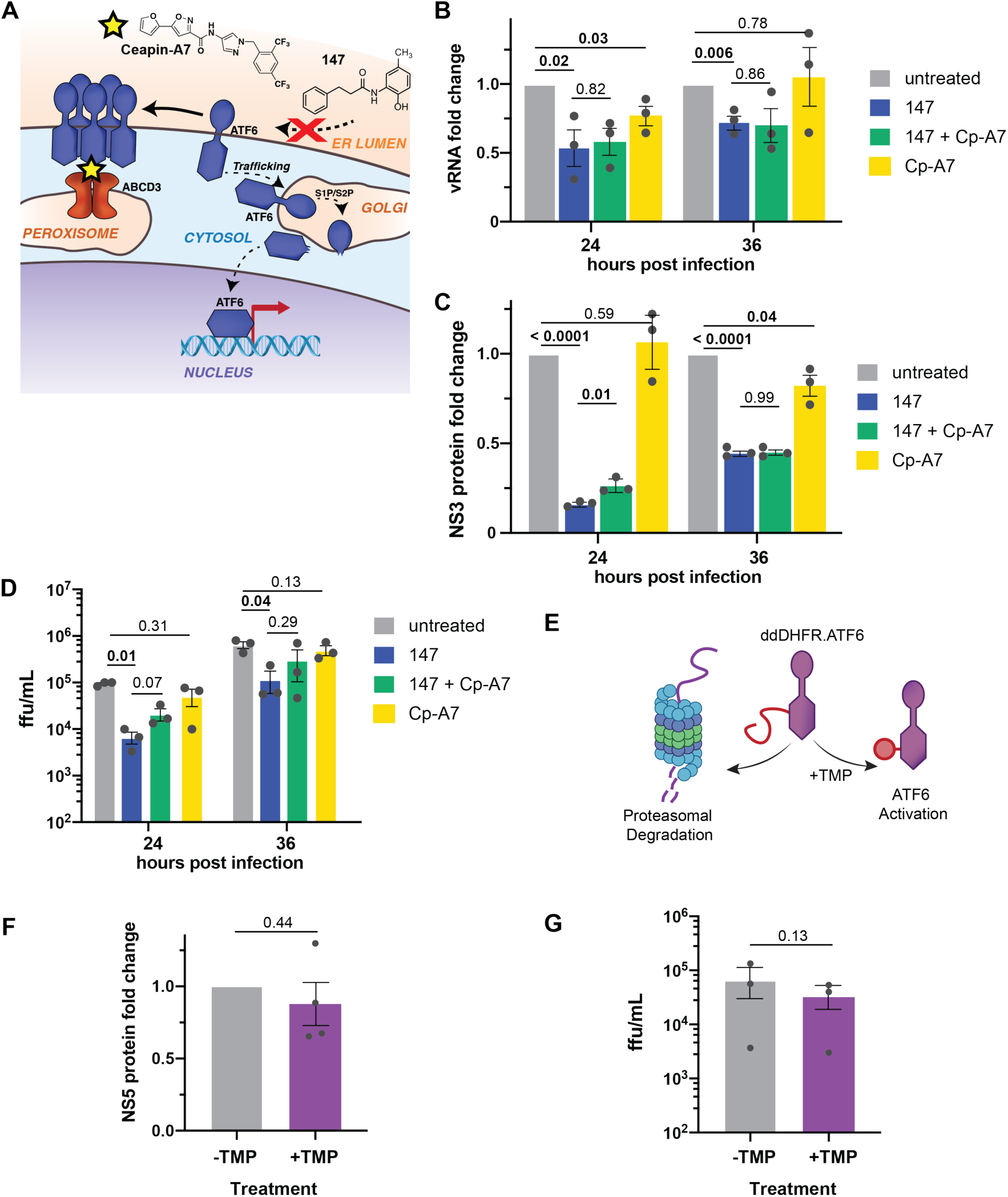
ATF6 inhibition does not attenuate the 147-mediated reduction in DENV replication. A. **Ceapin-A7** (**Cp7**) acts as an ATF6 inhibitor downstream of **147** by tethering inactive ATF6 to the peroxisomal membrane protein ABCD3 preventing ATF6 trafficking to the Golgi and activation through S1/S2 cleavage. B. Bar graph showing the reduction in vRNA levels in response to **147** and **Cp7** treatment measured by qPCR in Huh7 cells infected with DENV as outlined in **Fig. 2D. Cp7** does not attenuate the **147**-mediate reduction. Error bars correspond to SEM from 3 biological replicates and p-values from unpaired t-tests are shown. C. Bar graph showing the reduction in NS3 viral protein levels in response to **147** and **Cp7** treatment measured by Western blot in Huh7 cells infected with DENV as outlined in **Fig. 2D. Cp7** does not attenuate the **147**-mediate reduction. Error bars correspond to SEM from 3 biological replicates and p-values from unpaired t-tests are shown. D. Bar graph showing the reduction in DENV viral titers in response to **147** and **Cp7** treatment in Huh7 cells infected with DENV as outlined in **Fig. 2D. Cp7** only minimally attenuate the **147**-mediate reduction at 24 hpi. Error bars correspond to SEM from 3 biological replicates and p-values from paired ratio t-tests are shown. E. Schematic of the destabilized-domain (dd)DHFR.ATF6 construct, which is constitutively degraded in the absence of the stabilizing ligand trimethoprim (TMP). TMP addition leads to accumulation of ATF6 and transcriptional activation of ATF6 target genes. F. Graph showing NS5 viral protein levels in DENV infected Huh7 cells that were transiently transfected with ddDHFR.ATF6. ATF6 was activated through addition of TMP 16 hours prior to DENV infection (MOI 3). Data points were collected 24 hpi. Error bars correspond to SEM from 3 biological replicates and p-value from unpaired t-tests is shown. G. Graph showing DENV viral titers in infected Huh7 cells that were transiently transfected with ddDHFR.ATF6. ATF6 was activated through addition of TMP 16 hours prior to DENV infection (MOI 3) and DENV ffu were quantified 24 hpi. Error bars correspond to SEM from 3 biological replicates and p-value from ratio paired t-tests is shown.

To further probe whether the reduced viral propagation could be ascribed to activation of the ATF6 pathway, we took advantage of an orthogonal chemical genetic approach to selectively induce the ATF6 pathway independent of global ER stress. We transiently transfected a destabilized DHFR.ATF6 construct into Huh7 cells. This construct is constitutively degraded in the absence of a small molecule ligand, but can be stabilized through addition of trimethoprim (TMP) leading to accumulation of DHFR.ATF6 and selective induction of ATF6-regulated genes (**Fig. 3E**) (Shoulders et al., 2013). We confirmed TMP-dependent upregulation of ATF6-regulated targets BiP, PDIA4, and GRP94 in Huh7 cells (**Fig. S3B-C**). Next, we pre-treated Huh7 cells with TMP to activate DHFR.ATF6, infected cells with DENV2, and quantified propagation of the virus by monitoring viral protein levels by Western blot and measuring infectious titers. Chemical-genetic ATF6 activation did not lead to a measurable reduction in viral protein levels (**Fig. 3F**). At the same time, a moderate decrease in viral titers could be observed, but this reduction in viral propagation was much lower than seen with **147** treatment (**Fig. 3G**). Together, the results from the ATF6 inhibitor and chemical-genetic ATF6 activation indicate that induction of ATF6-regulated ER proteostasis fators does not impact vRNA replication or production of viral proteins, and only partially accounts for the reduction in DENV2 viral titers.

### Reduced DENV propagation requires 147 to covalently target protein thiols

Considering that reduced viral propagation in response to **147** was only partially mediated by ATF6 activation, we sought to explore other mechanisms of how the molecule could impair the virus. Previous studies showed that **147** is a prodrug and requires metabolic activation in cells to generate a reactive *p*-quinone methide that can then form protein adducts with reactive cysteine residues (**Fig. 4A**) (Palmer et al., 2020; Paxman et al., 2018; Plate et al., 2016). In several cell types, ER-resident protein disulfide isomerases (PDIs) were identified as the common protein targets of **147**, and this modulation of PDIs was linked to the activation of ATF6 (Paxman et al., 2018). To explore whether covalent targeting of reactive thiols by **147** is required for the reduction in DENV propagation, we blocked the covalent modifications through addition of an excess of the small-molecule thiol 2-mercaptoethanol (BME) to the cells treated with **147** (**Fig. 4A**). We confirmed that this addition did not impair cell viability (**Fig. S4B**), and that BME alone only had a minimal effect on viral protein levels (**Fig. S4A**). When BME was added to DENV infected cells that were treated with **147**, this resulted in a partial to complete recovery of NS5 protein at 24 and 36hpi, respectively (**Fig. 4B**). Similarly, the reduction in DENV2 viral titers was partially attenuated by the addition of BME (**Fig. 4C**). These results indicate that the targeting of cellular thiol groups is required for the inhibition of virus propagation.

**Figure 4 – (with 1 supplement).**
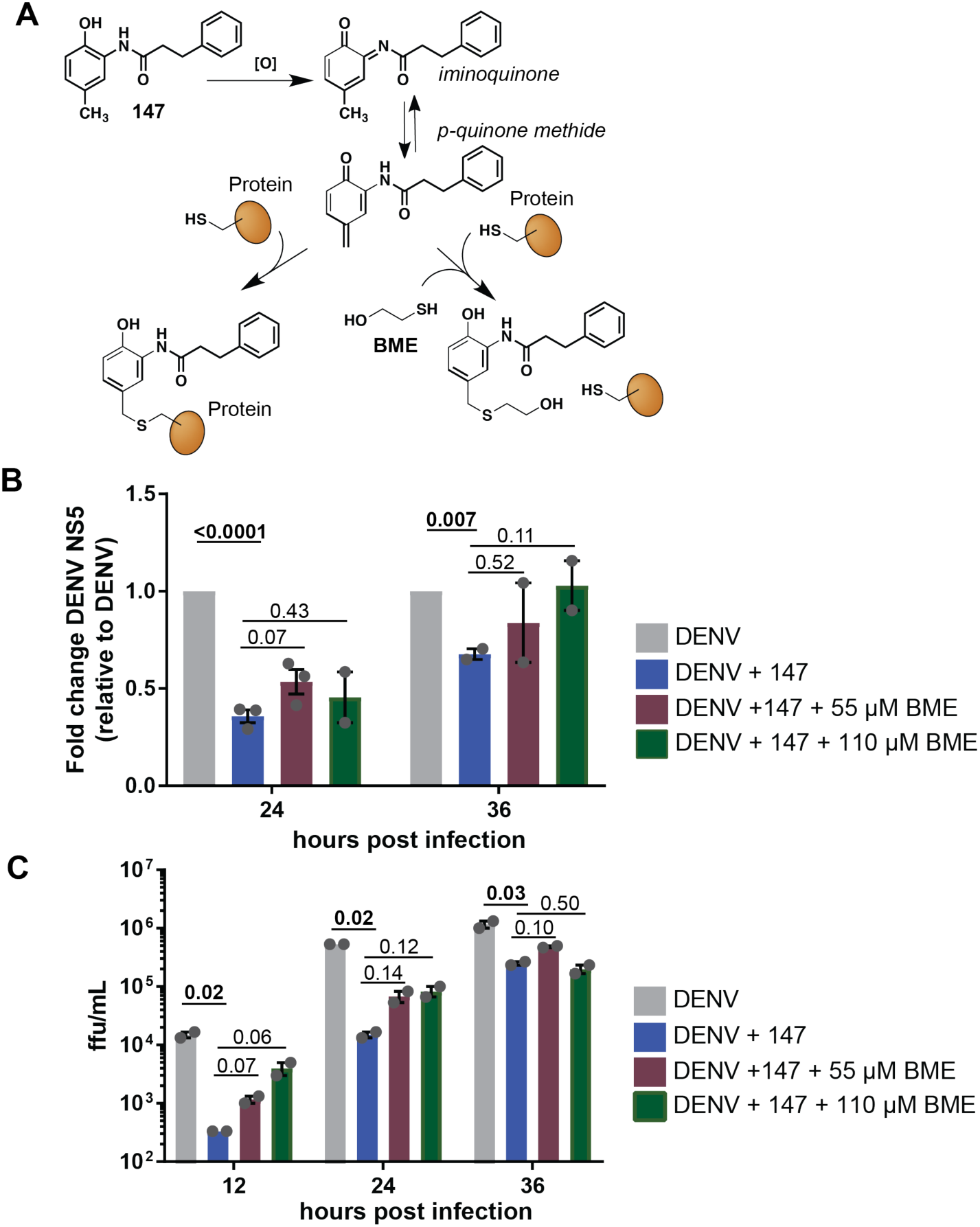
147-mediated reduction in DENV infection is sensitive to small molecule thiols. A. Schematic outlining the metabolic activation mechanisms of **147**. After oxidation by P450 enzymes, the generated *p*-quinone methide can react with thiol nucleophiles such as cysteine residues on cellular protein targets. Addition of exogenous free thiols such as 2-mercaptoethanol (BME) can quench the active form of **147** before it reacts with protein targets. B. Graph showing reduction in DENV NS3 and NS5 protein levels in response to **147** treatment and addition of BME. Huh7 cells were pretreated with **147** and or 55µM BME 16 hours prior to DENV infection (MOI 3). Protein levels were quantified by Western blot. Error bars show SEM and p-values from unpaired t-tests are shown. C. Bar graph showing DENV viral titer levels in response to 147 treatment and rescue through addition of BME. Huh7 cells were pretreated with **147** and or 55µM BME 16 hours prior to DENV infection (MOI 3) and viral ffu were quantified by a focus forming assay. Error bars show SEM and p-values from ratio paired t-tests are shown.

We next explored whether targeting of specific PDIs proteins by **147** was required for the reduced viral replication. Previous studies determined the covalent targets of **147** in HEK293T, HepG2, and ALLC plasma cells (Paxman et al., 2018). However, it was conceivable that the metabolic activation mechanism differs in cell types and could result in alternative targets. We therefore determined whether **147** could similarly target PDIs in Huh7 cells. We took advantage of the active analog **147-20**, which contains an alkyne handle that enables further Click chemistry derivatization with desthiobiotin after protein labeling, followed by isolation of the targeted proteins on streptavidin resin (**Fig. 5A**). The desthiobiotin probe also contained a TAMRA fluorophore that allowed gel-based visualization of the targeted proteins (**Fig. 5B**). We confirmed that **147-20** retained activity reducing DENV titer levels and NS3 protein levels in infected cells (**Fig. S5A-B**). In contrast, treatment with **147-4**, an inactive analog where generation of the quinone methide is blocked by the trifluoromethyl group, did not reduce infection. We probed for the presence of specific PDIs (PDIA4, PDIA1, PDIA6), which were identified as **147** targets in other cell lines. These proteins were clearly detectable at molecular weights of prominent labeled bands that were competed by **147**, confirming them as targets in Huh7 cells (**Fig. 5B, S5C**) (Paxman et al., 2018). In DENV-infected cells that were treated with **147-20**, no additional labeled protein bands were observed, indicating that viral proteins are not directly targeted by the compound (**Fig. S5D**).

**Figure 5 – (with 1 supplement).**
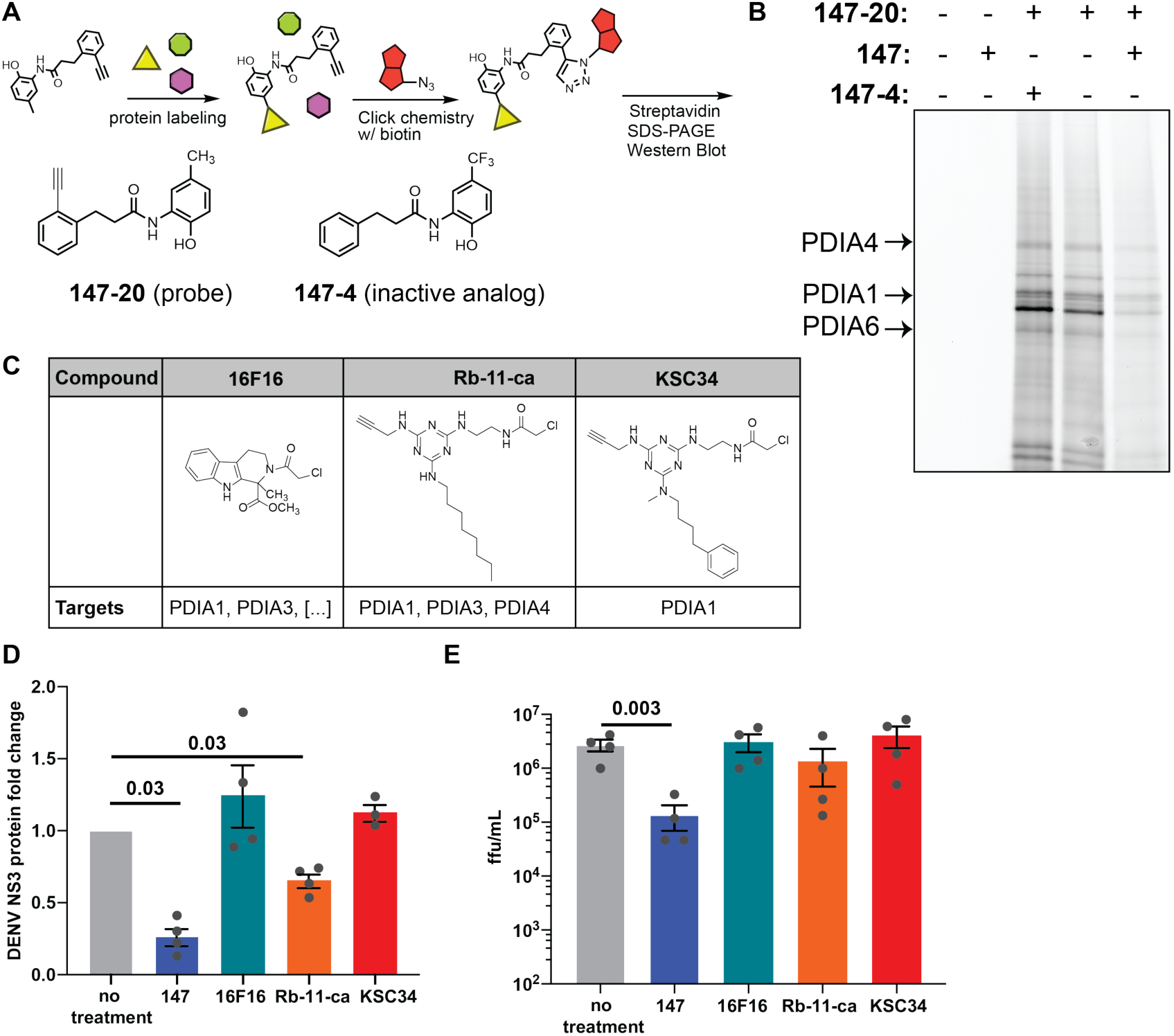
Modification of individual protein disulfide isomerases by 147 is not sufficient to reduce viral infection. A. Illustration of the chemoproteomic workflow for target identification using chemical derivatives of **147**. Cells are treated with **147-20**, an alkyne analog of 147 which retains activity and can covalently label proteins as outlined in **Fig. 4A**. The alkyne handle enables derivatization with either a fluorophore or biotin azide for detection or affinity purification of protein targets. **147-4** is an inactive analog of **147** used to determine specificity in competition experiments. B. Fluorescence SDS-PAGE image identifying proteins labeled by **147-20** in Huh7 cells. Huh7 cells were treated with **147-20** (3µM) and 3-fold excess competitor (active **147**, or inactive **147-4**) for 18 hours. Labeled proteins in cell lysates were derivatized with a TAMRA-desthiobiotin azide, proteins resolved on SDS-PAGE and a fluorescence image of the gel is shown. The pattern of labeled proteins reveals similar targets as seen in other cell lines (Paxman et al., 2018). The location of PDIA4, PDIA6, and PDIA1/P4HB is indicated. **Fig. S5C** shows images of Western blot overlays probing for individual PDI targets. C. Table showing chemical structures of PDI inhibitors **16F16, Rb-11-ca**, and **KSC-34** and their target selectivity for PDI isoforms. D. Graph quantifying DENV NS3 protein levels in Huh7 cells in response to treatment with small molecule PDI inhibitors. Huh7 cells were treated with **147** or PDI inhibitors 16 hours prior to DENV infection (MOI 3) and NS3 protein levels were quantified by Western blot 24 hpi. Only **147** and **Rb-11-ca** treatment lead to a significant reduction in viral protein. Error bars show SEM and p-values from unpaired t-tests are shown. E. Graph showing DENV viral titers in Huh7 cells treated with **147** or PDI inhibitors. Cells were treated with the corresponding molecules for 16 hours prior to DENV infection (MOI 3). Viral ffu were quantified 24 hpi by focus forming assay. Only **147** treatment reduces viral titers. Error bars show SEM and p-values from ratio paired t-tests are shown.

Considering the targeting of multiple PDI enzymes, we were interested in whether the inhibition of PDIs could have an important role in the reduction of viral propagation. We created stable knockdown cell lines of several PDIs using shRNAs (**Fig. S5E**). These cell lines were then infected with DENV2 and treated with **147** to assess the effect of PDI knockdown on propagation of the virus. Individual knockdown of *PDIA1* (*P4HB*), *PDIA4*, or *PDIA6* did not result in a decrease in viral titers indicating that none of these individual PDIs are essential for viral propagation (**Fig. S5F**). Only knockdown of *PDIA3* resulted in a small reduction in viral titers. Next, we measured viral infections in the knockout cell lines in the presence of **147** to determine whether particular PDIs are required for the **147**-mediated virus inhibition. We observed no measurable attenuation in the reduced viral titers with any single PDI knockdown, including the *PDIA3* knockdown, suggesting that no individual PDIs are responsible for the **147**-mediated effect (**Fig. S5F**).

To further explore whether PDI inhibition could be responsible for the reduction in DENV infection, we took advantage of several published small-molecule inhibitors of PDIA1 (**Fig. 5C**) (Banerjee et al., 2013; Cole et al., 2018; Hoffstrom et al., 2010). Importantly, these molecules display different degrees of specificity towards PDIA1 relative to other PDIs, mirroring the polypharmacology observed with **147**. We profiled the specific PDIs targeted by the compounds in Huh7 cells by modifying the alkyne handles on the molecule with a fluorophore. **KSC-34** displayed the greatest preference for PDIA1, while **Rb-11-ca** also labeled PDIA4, PDIA6, and several other proteins (**Fig. 5SG**). This is consistent with observations in other cell types and the development of **KSC-34** as a more specific derivative of **Rb-11-ca** (Cole et al., 2018). **16F16** does not contain an alkyne handle but previous studies show even broader reactivity towards PDIs than **Rb-11-ca** (Hoffstrom et al., 2010). We then tested the effect of the compounds on DENV2 infection in Huh7 cells using the same treatment regime as for **147. KSC-34** and **16F16** were unable to reduce viral NS3 protein levels (**Fig. 5D**). **Rb-11-ca** was able to lower viral protein production by 40%; however, this reduction did not translate into lower viral titers (**Fig. 5E**). Reduction in viral protein levels with **Rb-11-ca** could be attributed to compound toxicity, as we observed a 40% reduction in cell viability (**Fig. S5H**). These results indicate that targeting of individual PDIs is not solely responsible for the reduced viral titers by **147** and that these existing PDI inhibitors are not effective at inhibiting DENV replication.

To identify additional mechanisms that could contribute to the **147**-mediated reduction in DENV viral propagation, we turned to untargeted quantitative proteomics of the affinity-enriched samples to identify additional protein targets. We used tandem mass tags for relative quantification of proteins in the **147-20** enriched samples relative to the competition samples (**147-20** treated with 3-fold excess **147**) (**Fig. 6A**). This analysis confirmed that PDIs were the most highly enriched proteins that were covalently targeted by **147-20** (**Fig. 6B, Fig. S6A, Table S2**). While labeling of PDIA4, PDIA6, and PDIA1 was confirmed above, we identified additional targeted PDIs (TXNDC5, PDIA3, TXM1) and two other highly enriched protein targets (GSTO1 and ALDH1A1). We compared these targets to previous datasets of **147-20** targets in HEK293T, HepG2, and ALMC-2 plasma cells (**Fig. 6C, Fig. S6B**) (Paxman et al., 2018). The additional proteins were targets specific to Huh7 cells and were not observed in other cell types, suggesting that they could have a unique role in the reduction in virus propagation in the Huh7 cells.

**Figure 6 – (with 1 supplement).**
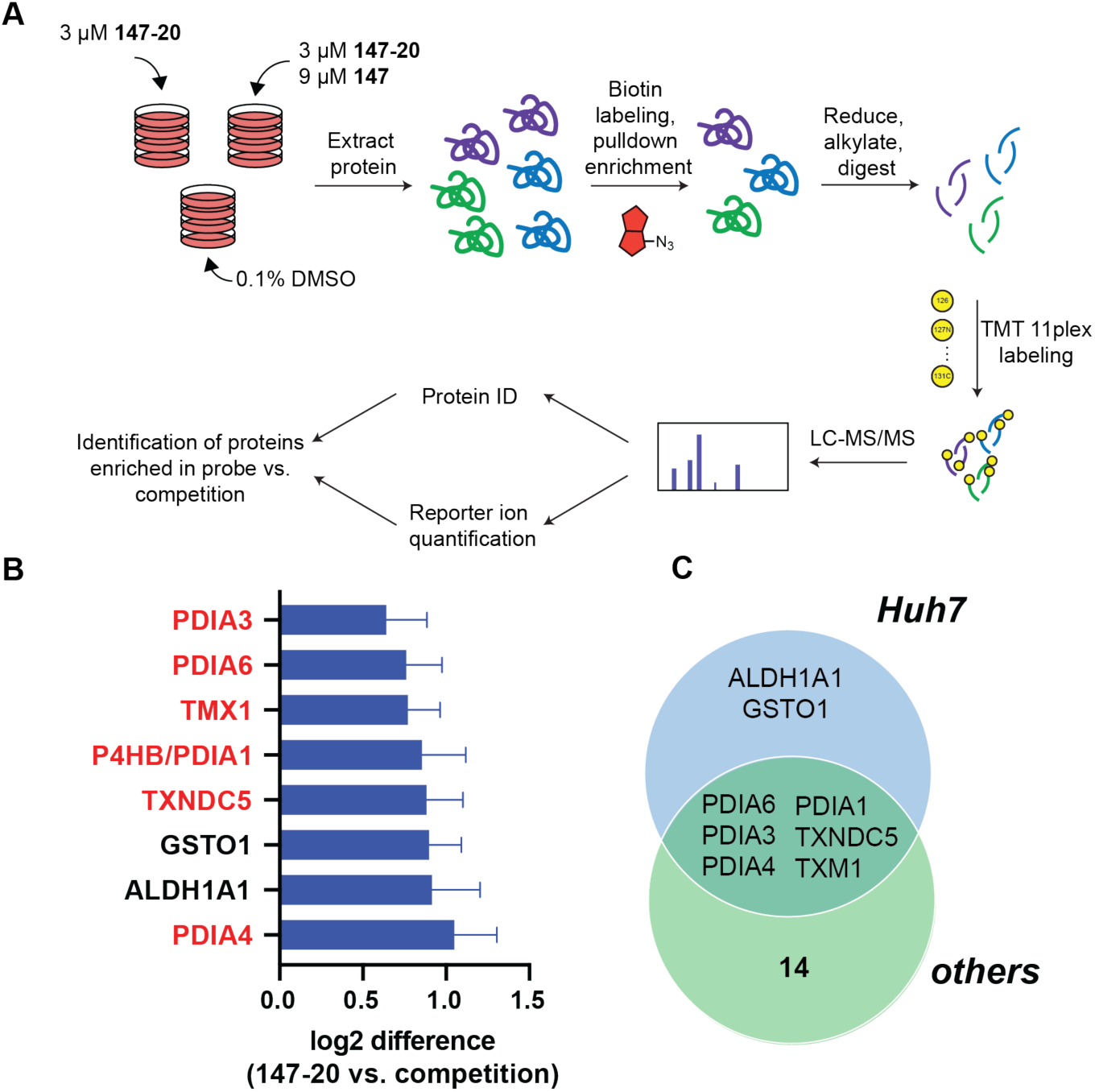
Identification of covalent protein targets of 147 in Huh7 cells. A. Illustration of the workflow for the chemoproteomic target identification. Huh7 cells were treated with 3µM **147-20** alone or in competition with 9µM **147** for 18 hours. Cell lysates were labeled with a TAMRA-desthiobiotin probe, labeled proteins were isolated on streptavidin beads, eluted and digested with trypsin. Individual samples were then labeled with tandem mass tags, pooled and subjected to LC-MS/MS for identification of proteins and quantitative comparison of proteins in the **147-20** treated compared to the **147-20**/**147** competition. B. Graph showing the most highly enriched proteins targets in the **147-20** treated samples compared to the competition with **147**. The full data is shown in **Fig. S6A**. High-confidence targets were filtered that displayed a log2 enrichment ratio greater than two standard deviations of the distribution of enrichment ratios across four biological replicates. C. Venn diagram showing the comparison of high-confidence protein targets of **147-20** in Huh7 cells and other cells lines (HEK293T, HepG2 and ALMC-2) identified in a prior study (Paxman et al, 2018). **Fig. S6B** shows the overlapping targets with individual cell lines.

### Compound 147 is effective against other DENV strains and Zika virus

Studies until this point were carried out with Dengue serotype 2 BID-V533 (isolated in 2005 in Nicaragua) in Huh7 cells. To determine the scope of **147** antiviral activity, we tested the effect against DENV-2 propagation in HepG2 cells, another liver carcinoma cell line commonly used as an infection model. Prior data also showed that **147** treatment is non-toxic in HepG2 cells and can reduce amyloidogenic protein secretion (Plate et al., 2016). When testing the impact on DENV titers in HepG2 cells, **147** treatment resulted in a significant reduction in DENV propagation 48 hpi (**Figure 7A**). A similar decrease in protein levels was also detected (**Fig. S7A**). Next, we proceeded to test whether the compound could be more broadly effective against other DENV strains. We tested **147** against a different DENV-2 strain (16681, isolated in 1984 in Thailand) as well as against strains from DENV serotypes 3 and 4 (Kinney et al., 1997). Treatment with compound **147** was able to reduce viral titers consistently greater than 95% (**Fig. 7B–D**) and protein levels were similarly reduced (**Fig. S7B**). Finally, we sought to test whether compound **147** could be useful to treat other flaviviruses. We measured the effect against two different strains of Zika virus; MR766 (isolated from Eastern Africa in the 1950s), and PRVABC59 (isolated from Puerto Rico in 2015) (Dick and Kitchen, 1952; Lanciotti et al., 2016). The molecule displayed similar activity at reducing ZIKV titers and protein levels for both strains (**Fig. 7E-F, Fig. S7C-D**). These results provide evidence that compound **147** is broadly active against flaviviruses in multiple cell lines, thus facilitating its potential application as an extensive host-centered antiviral agent.

**Figure 7 – (with 1 supplement).**
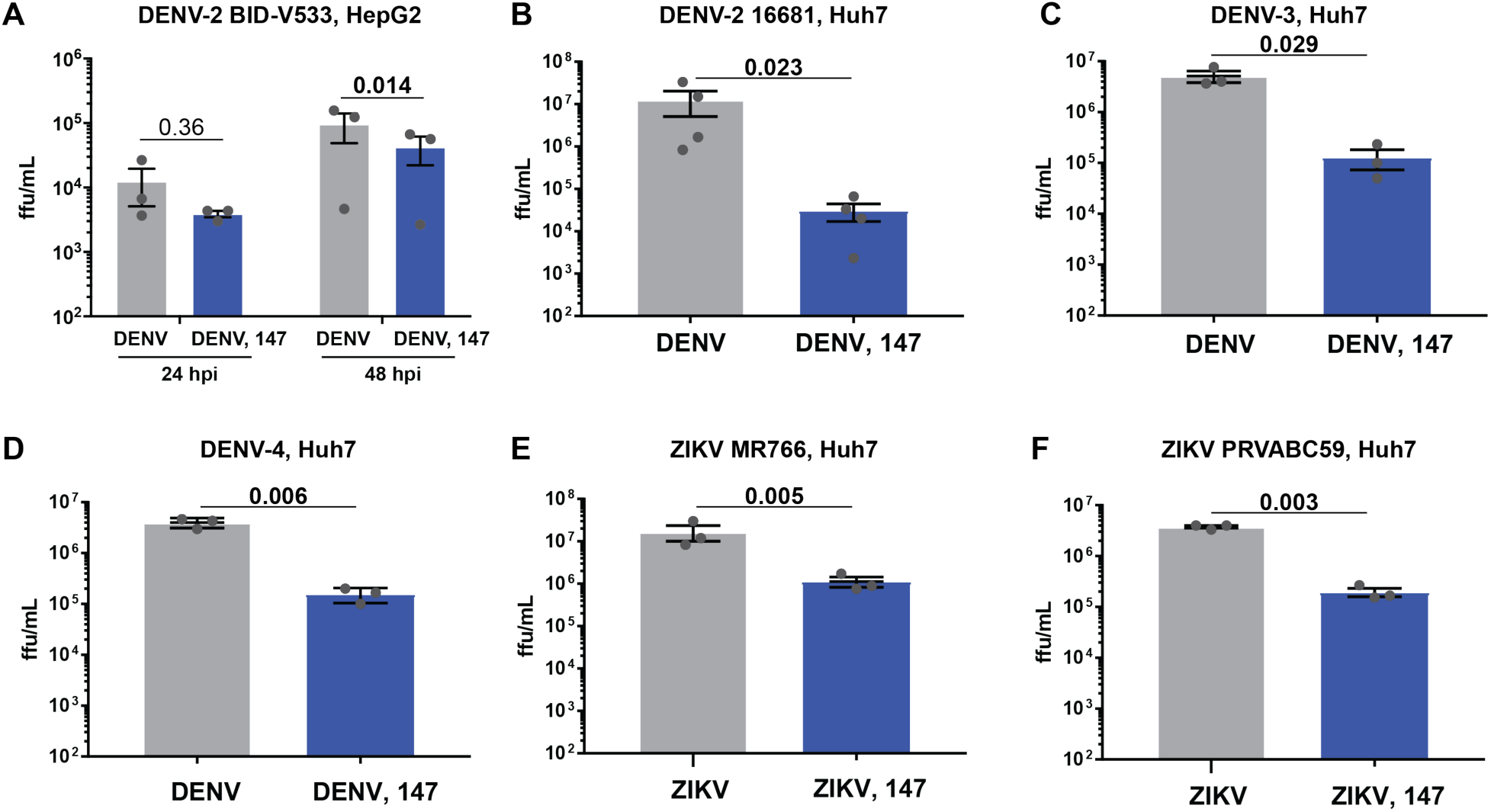
Compound 147 can reduce infection of multiple DENV strains, serotypes, as well as Zika virus. A. Reduction of DENV-2 infection in response to **147** in HepG2 liver carcinoma cells. HepG2 cells were pretreated with **147** (10µM) for 16 hours prior to infection with DENV-2 BID-V533 (MOI 3). Viral titers were determined 24 and 48 hpi by focus forming assays. B. Graph showing reduction in infection with DENV serotype 2 strain 16681 (isolated in Thailand in 1984) in response to treatment with compound **147**. Prior studies were carried out with DENV-2 BID-V533 (Nicaragua). Huh7 cells were pretreated with compound **147** (10µM) 16 hrs prior to DENV infection (MOI 3) and viral titers were determined 24 hpi. C.–D. Treatment with **147** reduces infection of DENV serotype 3 and 4. Huh7 were pretreated with **147** (10µM) for 16 hours prior to infection with DENV-3 Philippines/H87/1956 (**C**) or DENV-4 H241 (**D**) (MOI 3). Viral titers were determined 24 hpi by focus forming assays. E.–F. Compound **147** is similarly active at reducing infection of Zika virus. Huh7 cells were pretreated with **147** (10µM) for 16 hours prior to infection with ZIKV strain MR766 (**E**) or ZIKV strain PRVABC59 (**F**). Viral titers were determined 24 hours hpi by focus forming assay. All error bars show SEM from 3 biological replicates and p-values from ratio paired t-tests are shown.

## DISCUSSION

The continued health burden from arbovirus infections, such as Dengue and Zika, combined with the lack of effective vaccines and therapeutics against these viruses, highlights a need for the development of new antiviral strategies. Therapeutic methods which target host factors required for viral propagation provide a path unlikely to elicit drug resistance and may act broadly across several virus species (Aviner and Frydman, 2020; Geller et al., 2013, 2007). However, a challenge remains to target host factors that selectively impair viral replication without causing toxicity to the host cell and infected organism. Prior studies with **147** have already indicated that the compound has broad potential to safely ameliorate proteostasis imbalances that are associated a variety of disease conditions (Blackwood et al., 2019; Plate et al., 2016). The compound is effective at reducing the secretion and aggregation of amyloidogenic proteins, such as immunoglobulin light chains and transthyretin (Plate et al., 2016). In addition, selective remodeling of ER proteostasis pathways through an ATF6-dependent mechanism by **147** was shown to prevent oxidative organ damage in mouse models of ischemia/reperfusion (Blackwood et al., 2019). The same study also showed that **147** could be safely administered to mice through IV injections and was able to activate ER proteostasis pathways in multiple tissues, including liver, kidney and heart (Blackwood et al., 2019). Here, we further expand the therapeutic potential for compound **147** by establishing that the small molecule serves as a promising strategy to target multiple flaviviruses without causing significant toxicity in the host cell.

Treatment with the small molecule proteostasis regulator, **147**, resulted in a decrease in DENV vRNA, protein, and titer levels. The titer levels were the most affected, indicating that the compound was most active at preventing the assembly and secretion of mature infectious virions. This is consistent with the compound targeting critical proteostasis processes that could be required for production of infectious virions and are thus, crucial to the viral life cycle (Fischl and Bartenschlager, 2011; Heaton et al., 2016).

**147** was designed as a preferential activator of the ATF6 branch of the UPR. Given that DENV innately activates the ATF6 signaling pathway, it was unexpected that small molecule activation of ATF6 could impair virus propagation. However, addition of the ATF6 inhibitor **Cp7** was not effective at reducing virus replication, indicating that ATF6 activation did not mediate the antiviral effect. **Cp7** on its own also did not reduce virus infection, which is consistent with prior knockout of ATF6 not affecting DENV propagation (Peña and Harris, 2011). These results prompted us to examine alternative mechanisms for the antiviral effects by investigating the specific protein targets of **147**.

Prior work showed that **147** is a prodrug that is metabolically oxidized to an iminoquinone or quinone methide, which covalently targets nucleophilic cysteine residues on cellular protein (Palmer et al., 2020; Paxman et al., 2018; Plate et al., 2016). We showed that the antiviral activity of **147** could be attenuated with BME treatment, demonstrating that the metabolic activation mechanism and covalent targeting of reactive thiols is similarly required to reduce virus infection. We repeated the chemoproteomic target identification in Huh7 cells using an analog of **147** and identified a similar set of PDIs, including PDIA1, PDIA4, and PDIA6, as targets in this cell line. However, knockdown of individual PDIs was not sufficient to reduce viral titers suggesting that these individual PDIs are not responsible for the inhibition of viral infection. This raises the possibility that 1) polypharmacologic targeting of multiple PDIs may be necessary to sufficiently impair virus propagation, or 2) alternative **147**-targets are responsible for the antiviral effects.

We addressed whether inhibition of multiple PDIs could be required by testing the effect of existing PDI inhibitors that display varying selectivity towards PDIA1/P4HB and additional PDIs also modified by **147**. None of the PDI inhibitor compounds significantly reduced DENV viral titers, suggesting that **147** elicits its antiviral activity through an alternative mechanism of action. Importantly, one of the compounds, **Rb-11-ca**, exhibited considerably more toxicity in Huh7 cells, even at the highest concentration that we tested for **147**. Toxicity has also been observed for other PDI inhibitors (Langsjoen et al., 2017). This further highlights the unique properties of compound **147**. The low toxicity could be explained by previous observations showing that less than 25% of the PDIA4 pool is labeled by the compound, indicating that PDIs are not globally inhibited by the molecule (Paxman et al., 2018). Furthermore, endogenous protein secretion and formation of disulfide bonds in secreted proteins, including fully assembled immunoglobulins, is not affected by **147** (Plate et al., 2016). This lack of perturbation to normal proteostasis processes and selective inhibition of viral propagation makes **147** an ideal candidate for a host-centered antiviral strategy.

To address the second possibility that alternative protein targets besides the most prominent PDIs could be responsible for reduction in virus propagation, we compared our identified targets in Huh7 cells to the prior list of covalently modified proteins in other cell lines (Paxman et al., 2018). In addition to the PDIs studied above, additional members of the PDI family were identified (TXNDC5, TXM1), as well as two unique proteins that were identified only in Huh7 cells: ALDH1A1 and GSTO1. These new targets also have catalytic Cys residues that are likely modified by **147**. Ongoing studies are directed at determining the potential role of these protein targets in the **147**-mediated antiviral mechanism.

The recent COVID-19 pandemic has refocused attention to the need for broad-spectrum antiviral agents that could inhibit future virus threats (Carrasco-Hernandez et al., 2017). Targeting essential host pathways that are commonly hijacked by virus infection without causing significant host toxicity is recognized as one important strategy to combat future infection with unknown pathogens. We determined that compound **147** has broad antiviral activity against multiple DENV strains and serotypes and that the compound is also effective against multiple ZIKV strains. These results highlight broad utility of the proteostasis regulator compound to reduce flavivirus infection by targeting conserved host cell processes that are commonly exploited by the virus. Our results here now provide an entirely new therapeutic application of **147** as a broad antiviral agent to reduce flavivirus infections.

## MATERIALS AND METHODS

### Key resources table

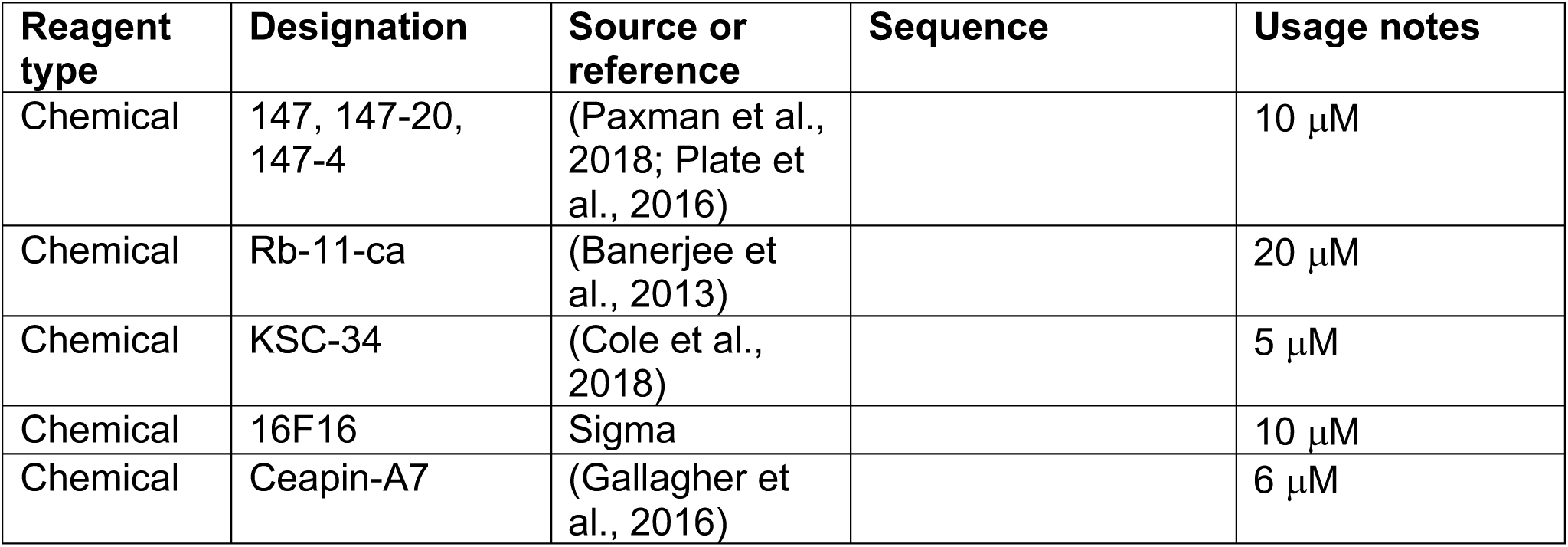
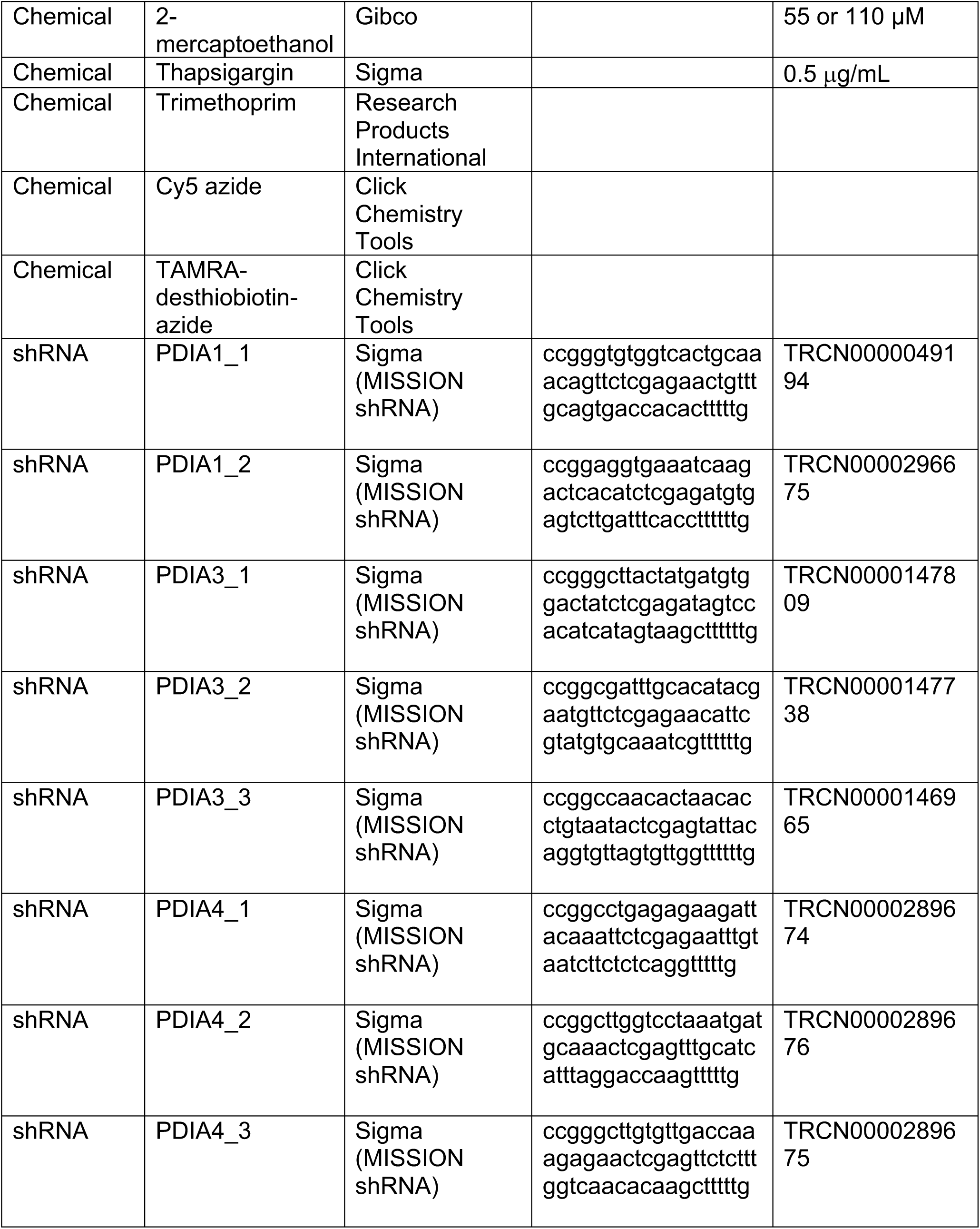
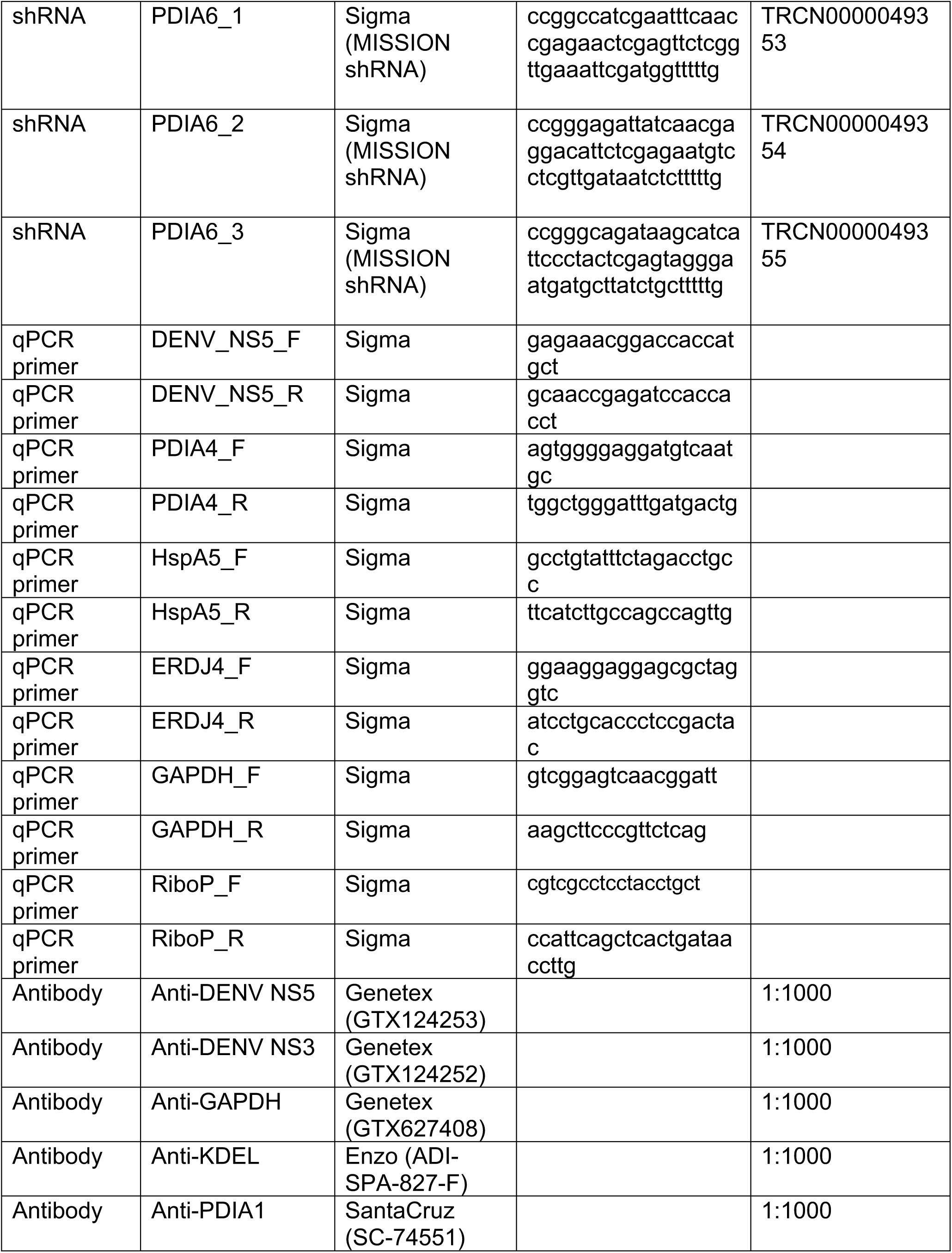
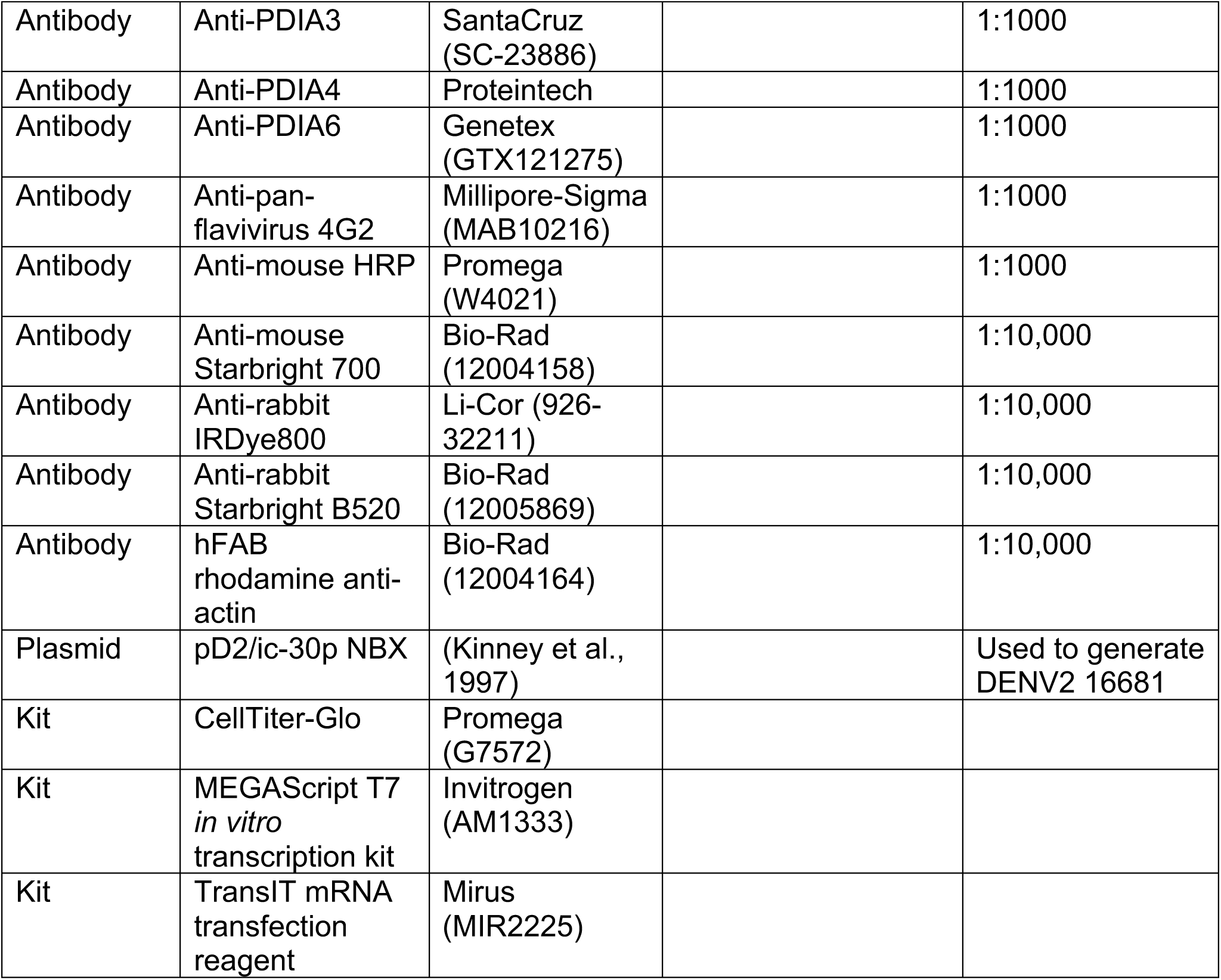

### Chemicals

All compounds except 2-mercaptoethanol (BME) were kept as 1000x stocks in DMSO (0.1% final concentration DMSO for cellular treatments). BME was directly diluted from a 55 mM DPBS stock. **147, 147-4, 147-20**, and **16F16** were used at 10 μM unless otherwise noted. **Rb-11-ca** was used at 20 μM. **KSC-34** was used at 5 μM. **Ceapin-A7** was used at 6 μM. Thapsigargin was used at 0.5 μg/mL.

### Cell Lines and Culture Conditions

Cells were maintained in Dulbecco’s Modified Eagle’s Medium (DMEM) with high glucose and supplemented with 10% fetal bovine serum (FBS), 1% penicillin/streptomycin, and 1% glutamine. All cell lines except C6/36 were kept at 37°C, 5% CO_2_. C6/36 cells were kept at 28°C, 5% CO_2_. Huh7 and HepG2 cells were obtained from ATCC. Vero and C6/36 cells were a kind gift from Dr. Tom Voss of the virology core at the Vanderbilt Vaccine Center. Cells were tested monthly for mycoplasma contamination.

### Viruses

#### DENV

Strain BID-V533 was a generous gift from Dr. Tom Voss of the Vanderbilt Vaccine Center. Strain 16681 was generated from a cDNA clone in pD2/ic-30p NBX that was a generous gift from Dr. Claire Huang at the Center for Disease Control. DENV-3 and DENV-4 strains were obtained from BEI Resources. For passaging, C6/36 or Vero cells were infected at MOI 0.1. Six days (C6/36) or three days (Vero) post-infection, culture supernatant was harvested and cell debris was pelleted at 3700xg for 10 minutes. Cleared supernatant was combined with 23% FBS and mixture was frozen as 1 mL aliquots at -80°C.

For experimental infections of cell lines, media was removed from cells and virus was added at MOI 3 for 3 hours (unless otherwise noted). Inoculum was removed, cells were washed, and media containing compounds or DMSO was added for the remainder of the experiment. Strain 16681 was generated from a cDNA clone (Kinney et al., 1997). Plasmid DNA was transformed into DH5α cells and purified by miniprep (Zymo Research). DNA was ethanol/sodium acetate precipitated for purification and concentration. Plasmid DNA was digested using XbaI (NEB) to cut upstream of the T7 promoter. A MEGAScript T7 in vitro transcription kit (Invitrogen) was used to obtain viral RNA, which was then transfected into Huh7 cells using the TransIT mRNA transfection reagent (Mirus). Virus stocks were amplified as indicated above.

#### ZIKV

Strains MR766 and PRVABC59 were obtained from ATCC. For passaging, Vero cells were infected at MOI 0.05. Three days post-infection, culture supernatant was harvested and cell debris was pelleted at 17000xg for 10 minutes. Cleared supernatant was used as virus stock and stored at -80°C. For experimental infections of cell lines, media was removed from cells and virus was added at MOI 0.5 for 1 hour. Inoculum was removed, cells were washed, and media containing compounds or DMSO was added for the remainder of the experiment.

#### Lentivirus

The 3^rd^ generation system was used for production. Respective shRNA plasmids were transfected into HEK293T cells together with lentivirus packaging plasmids RRG, VSVG, and Rev using the calcium phosphate method as described (Paxman et al., 2018). Media was changed 20 hours post-transfection. Supernatants were collected 3 days post transfection, and cleared at 3700xg for 10 minutes. To generate respective shRNA stable knockdown cell lines in Huh7 cells, lentiviruses prepared with pools of 2-3 shRNAs per target were added to Huh7 cells and media containing 4 μg/mL final concentration polybrene. Selection proceeded for 1 week under puromycin (5 μg/mL). Successful knockdown was confirmed by Western Blot.

### Viral Focus Forming Assay

Confluent Vero cells in 96 well plates were inoculated with 10-fold serial dilutions of DENV or ZIKV in BA-1 diluent (1xM199 media, 5% BSA, 1x L-glutamine, 1x penicillin/streptomycin, 0.04% sodium bicarbonate, 50 mM Tris) for 2 hours. The cells were overlaid with a 1:1 mixture of 2x nutrient overlay (2x Earle’s Balanced Salt Solution, 2x Ye-Lah Medium, 4% FBS, 0.4% sodium bicarbonate, 0.1 mg/mL gentamycin, 0.5 mg/mL amphotericin B) + 2.4% methylcellulose in water. After 2 days (ZIKV, DENV3, DENV4) or 3 days (DENV2), overlay was removed, and cells were fixed in ice cold 85% acetone for 30 minutes. Infectious foci were stained with a primary pan-flavivirus 4G2 antibody (1:1000 in 5% BSA, TBST) and secondary HRP antibody (1:1000 in 5% milk, TBST), then visualized using 400 μL 8 mg/mL 3-amino-9-ethylcarbazole (Sigma) in 10 mL 50 mM sodium acetate (Fisher) + 4 μL 30% H_2_O_2_ (Fisher) and exposed for 15-45 minutes until foci were visible.

### SDS-PAGE Gels and Immunoblotting

Cell pellets were lysed in RIPA buffer (50 mM Tris pH 7.5, 150 mM NaCl, 0.1% SDS, 1% Triton-X-100, 0.5% sodium deoxycholate) + Roche cOmplete protease inhibitor. Cells were left on ice for at least 10 minutes and lysate was cleared at 17,000xg for 10 minutes. Cleared lysate concentrations were normalized and gel samples were separated by SDS-PAGE. Gels were transferred to PVDF membranes using standard settings on the TransBlot Turbo (BioRad). Blots were blocked in 5% non-fat dry milk in Tris-buffered saline (TBST, 50mM Tris pH 7,4, 150mM NaCl, 0.1 % Tween-20) for 30 minutes before applying antibodies. Primary antibodies (in 5% BSA, TBST) were applied at room temperature for 2 hours or at 4°C overnight. Secondary antibodies (in 5% milk, TBST) were applied at room temperature for 30 minutes or at 4°C for 1 hour. All primary antibodies plus the anti-mouse HRP secondary were used at a 1:1000 dilution; all secondary antibodies were used at a 1:10,000 dilution.

### Quantitative RT-PCR

RNA was prepared from cell pellets using the Zymo Quick-RNA miniprep kit. cDNA was synthesized from 500 ng total cellular RNA using random primers (IDT), oligo-dT primers (IDT), and Promega M-MLV reverse transcriptase. qPCR analysis was performed using Bio-Rad iTaq Universal SYBR Green Supermix combined with primers for genes of interest (listed below) and reaction were run in 96-well plates on a Bio-Rad CFX qPCR instrument. Conditions used for amplification were 95°C, 2 minutes, 45 repeats of 95°C, 10s and 60°C, 30s. A melting curve was generated in 0.5 °C intervals from 65 °C to 95 °C. C_q_ values were calculated by the BioRad CFX Maestro software. Transcripts were normalized to a housekeeping gene (either RiboP or GAPDH). All measurements were performed in technical duplicate; each of these duplicates was treated as a single measurement for the final average. Data was analyzed using the BioRad CFX Maestro software.

### Cell Viability Assays

Cell viability was determined using the CellTiter-Glo reagent (Promega). Huh7 cells were seeded into 96 well plates into 100 μL culture media. 100x dilutions of compounds were prepared in culture media. 11 μL of each compound dilution was added to wells in triplicate. 16 hours after treatment, media was removed and cells were infected as described above (for -DENV samples, no virus was added). Media was replaced 24 hours after infection and 100 μL Cell Titer Glo reagent was added. Plates were gently shaken for 10 minutes, then luminescence was measured for each well on a Synergy HT plate reader. Luminescence values for each condition and/or compound treatment were averaged and normalized to a DMSO control. Where multiple independent replicates are indicated in legends, samples were averaged, and standard error of the mean was propagated.

### Enrichment of Covalent Protein Targets

Huh7 cells were treated with 3 μM **147-20** or a combination of 3 μM **147-20** and 9 μM **147**. Cells were harvested 16-18 hours post-treatment and lysed as described above. Click chemistry using 100 μM Cy5 azide or TAMRA desthiobiotin azide was performed with 1.6 mM BTTAA, 0.8 mM copper sulfate, and 5 mM sodium ascorbate for 1 hour at 37°C, shaking at 500 rpm. All reagents were purchased from Click Chemistry Tools. For detection, samples from Cy5 reactions were run directly on SDS-PAGE gels for analysis. For enrichment of proteins, excess reagents were removed via methanol/chloroform precipitation; protein was washed twice with methanol before resuspension in 6 M urea. Samples were diluted in PBS and added to pre-washed (in PBS) streptavidin agarose beads (Thermo Scientific). Pulldowns were rotated for 1 hour at room temperature. Supernatant was removed and beads were washed six times in PBS and 1% SDS. Samples were eluted two times by incubation with 50 mM biotin, pH 7.2, in 1% SDS (PBS) for 10 minutes.

### Mass Spectrometry Sample Preparation

Protein lysate from DENV infected and non-infected Huh7 cells (10 µg of protein as measured using Bio-Rad protein assay dye reagent) was aliquoted and purified using methanol/chloroform precipitation. Streptavidin-enriched samples for target identification were used without normalization and purified by methanol/chloroform precipitation. Mass spectrometry samples were prepared as described previously (Blackwood et al., 2019). Briefly, protein was resuspended in 3 μL 1% Rapigest (Waters) and diluted with water and 0.5 M HEPES. Proteins were reduced with 5 mM TCEP (Thermo Scientific) and subsequently alkylated with 10 mM iodoacetamide (Sigma). Proteins were digested in sequencing-grade trypsin (Thermo Scientific) overnight at 37°C while shaking at 750 rpm. Peptides were labeled for 1 hour at room temperature using 11-plex tandem mass tag (TMT) reagents (Thermo Scientific) resuspended in 40% acetonitrile. Labeling reactions were quenched with 0.4% ammonium bicarbonate for 1 hour at room temperature. Samples were combined and acidified to pH 2 by addition of 5% formic acid. Pooled samples were concentrated using a SpeedVac and resuspended in 94.9% water/5% acetonitrile/0.1% formic acid. Rapigest cleavage products were pelleted by centrifugation at 14,000xg for 30 minutes.

### Mass Spectrometry Data Acquisition

Digested peptides (20 µg) were loaded onto a triphasic column filled sequentially with 1.5 or 2.5 cm layers of 5 µm 100 Å C18 resin (Phenomenex), Luna 5 µm 100 Å strong cation exchange resin (Phenomenex) and a second layer of C18 resin using a high pressure chamber(Fonslow et al., 2012). The column was washed for 30 minutes in 95% water/5% acetonitrile/0.1% formic acid (buffer A) and attached to an Ultimate 3000 nano LC system connected to a 20 cm, 100 µm ID fused-silica microcapillary column with a laser-pulled tip filled with 3 µm 100 Å C18 resin (Phenomenex). Mass spectrometry was carried out on a Q-Exactive HF or an Exploris 480 instrument (ThermoFisher). For whole cell lysate samples, multidimensional peptide identification technology (MuDPIT) analysis was performed with 10 µL sequential injections of 0 - 100% buffer C (500 mM ammonium acetate, diluted in buffer A), each injection increasing by 10%, followed by an injection of 90% buffer C/10% buffer B (100% acetonitrile/1% formic acid). For target ID samples, only the 0, 10, 20, 40, 60, 80, 100 and 90%C/10%B injections were carried out. Peptides were separated on a 90 min gradient from 5 % to 40% buffer B at a flow rate of 500nL/min, followed by a 5 min ramp to 60 - 80% buffer B. Electrospray ionization was performed from the tip of the microcapillary column at a voltage of 2.2 kV with an ion transfer tube temperature of 275° C. Data-dependent mass spectra were collected by performing a full scan from 300 – 1800 m/z (Q-Exactive HF) or 375 – 1500 (Exploris 480) at a resolution of 120,000 and AGC target of 1e6. Tandem mass spectrometry was performed using TopN (15 cycles, Q-Exactive) or TopSpeed (3 sec, Exploris 480) from each full scan using HCD collision energy of NCE 38 (Q-Exactive HF) or NCE 36 (Exploris 480), isolation window of 0.7 (Q-Exactive HF) or 0.4 (Exploris 480), with automatic maximum injection time, a resolution of 45,000 (Q-Exactive HF) or 30,000 + TurboTMT setting (Exploris 480), fixed first mass of 110 m/z, and dynamic exclusion of 10 s (Q-Exactive HF) or 45 s (Exploris 480).

### Mass Spectrometry Data Analysis

Peptide and protein identification, and TMT reporter ion quantification was analyzed using Proteome Discover 2.4. Searches were carried out with the SEQUEST node using a human proteome database (UniProt) with DENV proteins added manually and the following parameters: 20ppm precursor mass tolerance, minimum peptide length of 6 amino acids, trypsin cleavage with a maximum of 2 missed cleavages, static modifications of +57.0215Da (carbamidomethylation at C) and +229.1629 (TMT6/11plex), and dynamic modification of +15.995 (oxidation at M), N-terminal methionine loss (−131.040), and N-terminal acetylation (+42.011). Search results were filtered with Percolator using a decoy database of reversed sequences with a peptide false discovery rate of 1% and a minimum of 2 peptides for protein identification. TMT intensities were quantified using the reporter ion quantification node with TMT intensities from each channel being normalized using the total peptide amount, quantitative value correction enabled to correct for TMT impurities, and co-isolation threshold set to 25%. Intensities for each protein were calculated by summing the intensities of each peptide. A reference TMT channel of pooled samples was included for scaling of protein abundances across multiple mass spectrometry runs. For Target ID samples, Proteome Discoverer searches were identical to the whole cell lysate searches, except the normalization of TMT intensities was omitted in the reporter ion quantification node. Log2 transformations were performed on all TMT reporter ion intensities, and averages were determined for four **147-20** replicates and the four **147-20/147** competition replicates. Fold enrichment was determined as the log2 fold difference of the two averages. The log2 fold enrichment was fitted to a gaussian distribution and targets were defined as proteins with log2 fold difference greater than two standard deviations.

## ACKNOWLEDGEMENTS

We thank Dr. Tom Voss (Vanderbilt University Medical Center), Dr. Claire Huang (Center for Disease Control) and Dr. Eranthie Weerapana (Boston College) for providing critical reagents and material. We are grateful to Dr. Renã Robinson (Vanderbilt University) and members of her group for access to mass spectrometry instrumentation. We thank members of the Plate group for critical reading of the manuscript. K.M.A. was supported by T32 AI112541, and an NSF GRFP fellowship. J.P.D and S.M.L were supported by T32 GM008554. We thank Vanderbilt University for providing critical startup funds.

## AUTHOR CONTRIBUTIONS

L.P.: Conceptualization, Project administration, Supervision, Investigation, Formal analysis, Data curation, Writing-original draft, Writing—review and editing

K.M.A.: Supervision, Formal analysis, Data curation, Investigation, Writing-original draft, Writing—review and editing

J.P.D., S.M.L., R.T., S.C.T.: Formal analysis, Data curation, Investigation, Writing— review and editing

---

**Figure 1 – figure supplement 1.**
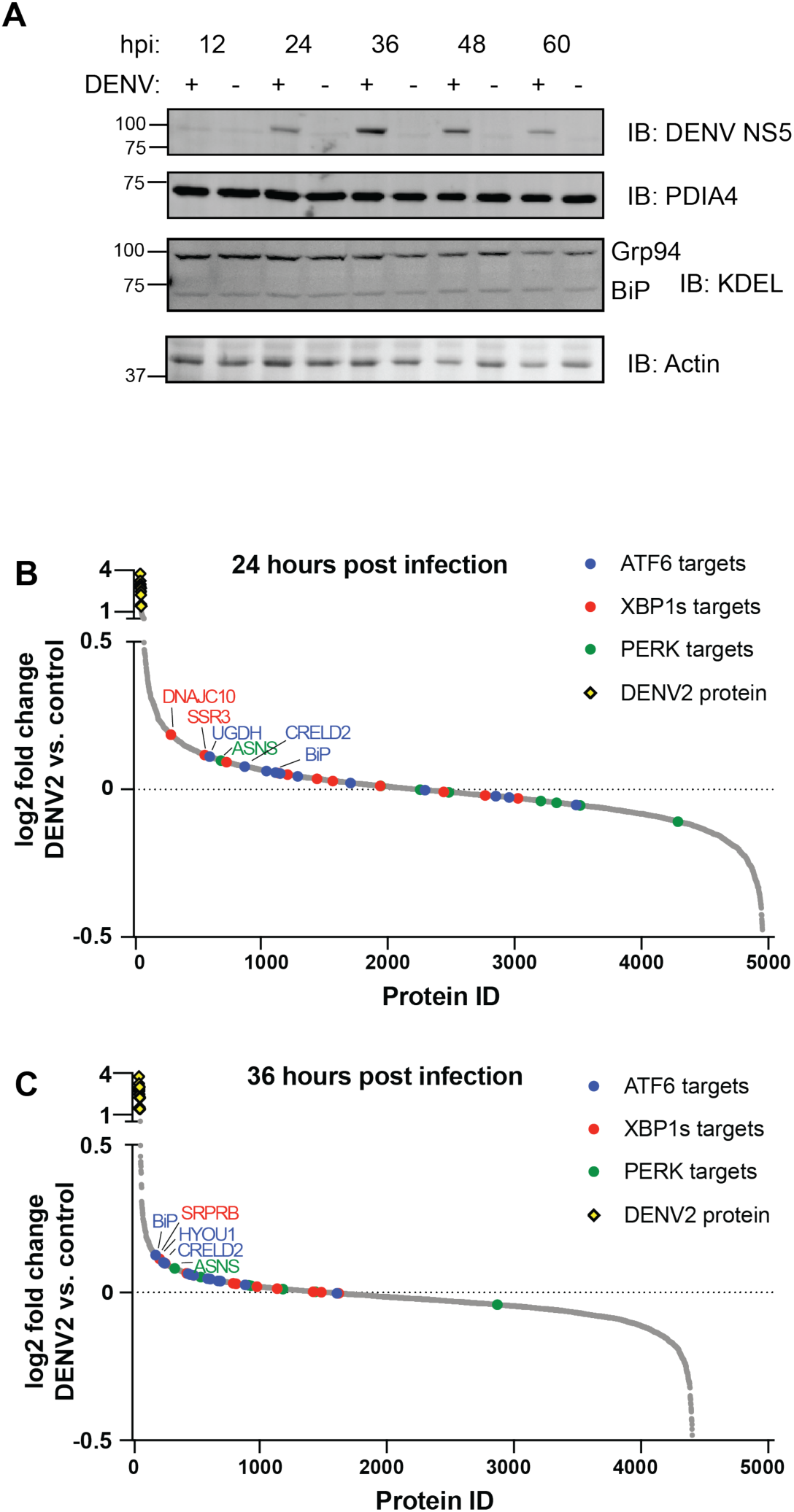
DENV infection induces the UPR. A. Western Blot showing time course of ATF6 upregulation over the course of DENV infection. Cells were infected with DENV-2 strain BID-V533 at MOI of 3 for 3 hours. Media was replaced and samples were taken at indicated timepoints post-infection. Cells were lysed, and protein samples were run on SDS-PAGE gels and visualized using PVDF membranes. Actin is included as a loading control. B.-C. Waterfall plots of proteomics data from samples infected with DENV-2 for 24 hours (**B**.) and 36 hours (**C**.), showing the presence of DENV proteins and upregulation of IRE1/XBP1s, ATF6, and PERK branches of the UPR. Cells were infected with DENV-2 strain BID-V533 at a MOI of 3 for 3 hours. Media was replaced and samples were collected at 24 or 36 hpi. Gene upregulation was quantified using TMT11plex reagents and comparison is between DENV-infected cells vs. noninfected cells. Graph represents 4-7 independent biological replicates. Source data is included in **Table S1**.

**Figure 2 – figure supplement 1.**
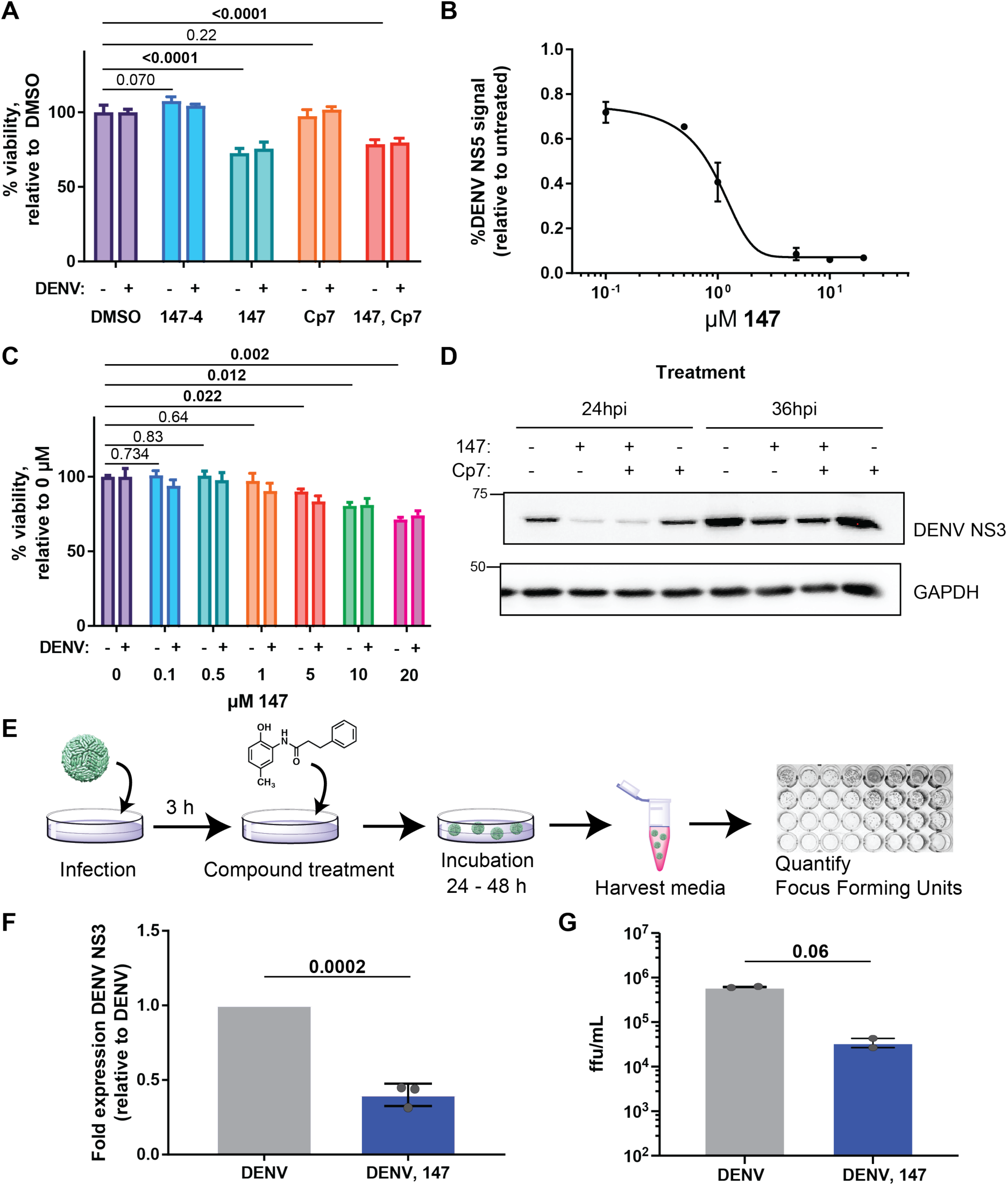
Treatment with small molecule 147 reduces DENV infection in a dose-dependent manner and only minimal impairs cell viability. A. Cell viability data showing **147** only modestly affects Huh7 cell viability over the course of treatment. No decrease in viability was seen with 147-4 or **Ceapin-A7** (**Cp7**). Cells were pretreated with 10 μM 147, 6 μM **Cp7**, or both for 16h, then infected (where indicated) with DENV-2 strain BID-V533 at a MOI of 3 for 3 hours. Media and treatments were replaced, and viability was measured 24hpi using the Promega Cell Titer Glo reagent. Graphs represent six independent biological replicates across two days of measurement. Error bars represent SEM. B. A dose-response curve shows DENV is sensitive to **147** treatment with an IC50 of approximately 1 μM. Cells were treated with the indicated concentrations of **147** for 16 hours, then infected with DENV-2 strain BID-V533 at a MOI of 3 for 3 hours. Media and treatments were replaced, and cells were collected 24 hpi. Cells were lysed, and protein samples were run on SDS-PAGE gels and visualized using PVDF membranes. Western blot intensities were normalized to a GAPDH loading control. Graphs represent 2-4 independent biological replicates. Error bars represent SEM. C. Increasing **147** concentration causes decreasing cell viability. No decrease in viability was seen with **147-4** or Ceapin-A7 (Cp7). Cells were pretreated with the indicated concentrations of **147** for 16h, then infected (where indicated) with DENV-2 strain BID-V533 at a MOI of 3 for 3 hours. Media and treatments were replaced, and viability was measured 24 hpi using the Promega Cell Titer Glo reagent. Graphs represent three biological replicates. Error bars represent SEM. D. Representative western blot showing **147** attenuates DENV protein levels. Cells were pretreated with 10 μM **147**, 6 μM **Cp7**, or both for 16h, then infected with DENV-2 strain BID-V533 at MOI of 3 for 3 hours. Media and treatments were replaced, and cells were collected at indicated timepoints. Cells were lysed, and protein samples were run on SDS-PAGE gels and visualized using PVDF membranes. GAPDH is included as a loading control. No visualization of DENV protein could be seen at 12 hpi. E. Eliminating the pretreatment step will determine if **147** blocks viral entry. Cells are only treated post-infection with 10 μM **147**, and focus forming units are quantified 24 hpi. F. Quantification of western blots with treatment only post-entry shows similar phenotype to treatment pre- and post-entry. Cells were only treated post-infection with 10 μM **147**, and cells were harvested 24 hpi. Cells were lysed, and protein samples were run on SDS-PAGE gels and visualized using PVDF membranes. Normalization to GAPDH was included as a loading control. Graph represents three independent biological replicates. Error bars represent SEM and p-value from an unpaired t-test is shown. G. Graph showing reduction in viral titers for only post-entry treatment. Cells were treated as in **E-F** and viral titers were determined by focus forming assay. Error bars represent SEM and p-value from a ratio paired t-test is shown.

**Figure 3 – supplement 1.**
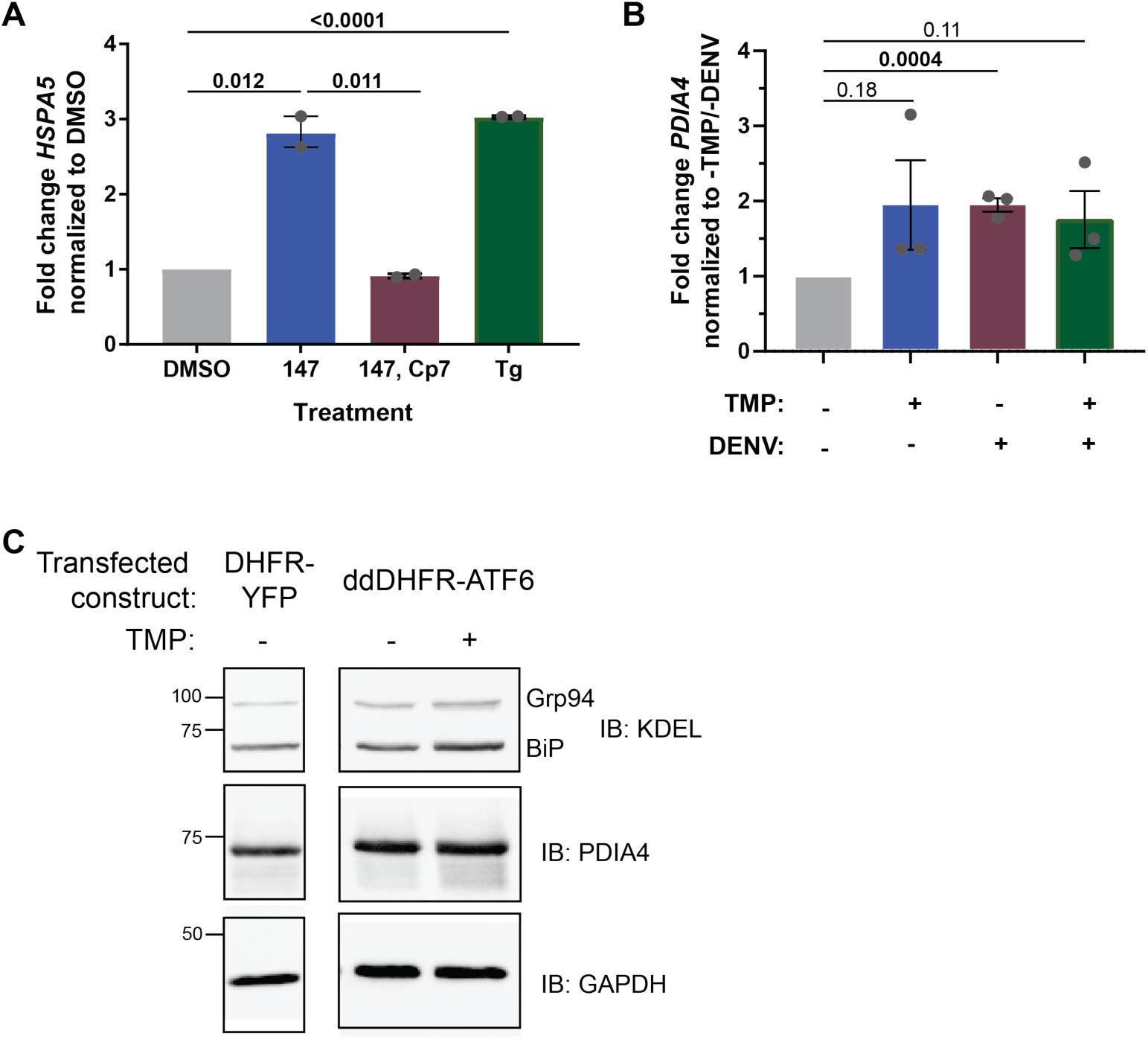
Ceapin-A7 blocks the 147-mediated ATF6 activation and ligand regulated DHFR.ATF6 activation in Huh7 cells. A. qPCR data showing **147** upregulates ATF6 target gene *HspA5/BiP*. Upregulation is reversed after cotreatment with **Cp7**. Thapsigargin (Tg), a global UPR stressor, is included as a positive control. Graphs represent two independent biological replicates, where each biological replicate is the average of two technical replicates. Error bars represent SEM and p-values from unpaired t-tests are shown. B. qPCR shows upregulation of ATF6 target gene *PDIA4* after transient transfection of ddDHFR-ATF6 construct AND treatment with trimethoprim (TMP). Treatment with DENV also shows slight upregulation of *PDIA4*. Combination TMP treatment and DENV infection does not appear to have a synergistic effect on *PDIA4* upregulation. Graphs represent 3 independent biological replicates, where each biological replicate is the average of two technical replicates. Error bars represent SEM and p-values from unpaired t-tests are shown. C. Western blot shows upregulation of ATF6 targets BiP, Grp94, and PDIA4 after transfection of ddDHFR-ATF6 construct AND treatment with TMP. A DHFR-YFP construct was used as a negative control.

**Figure 4 – supplement 1.**
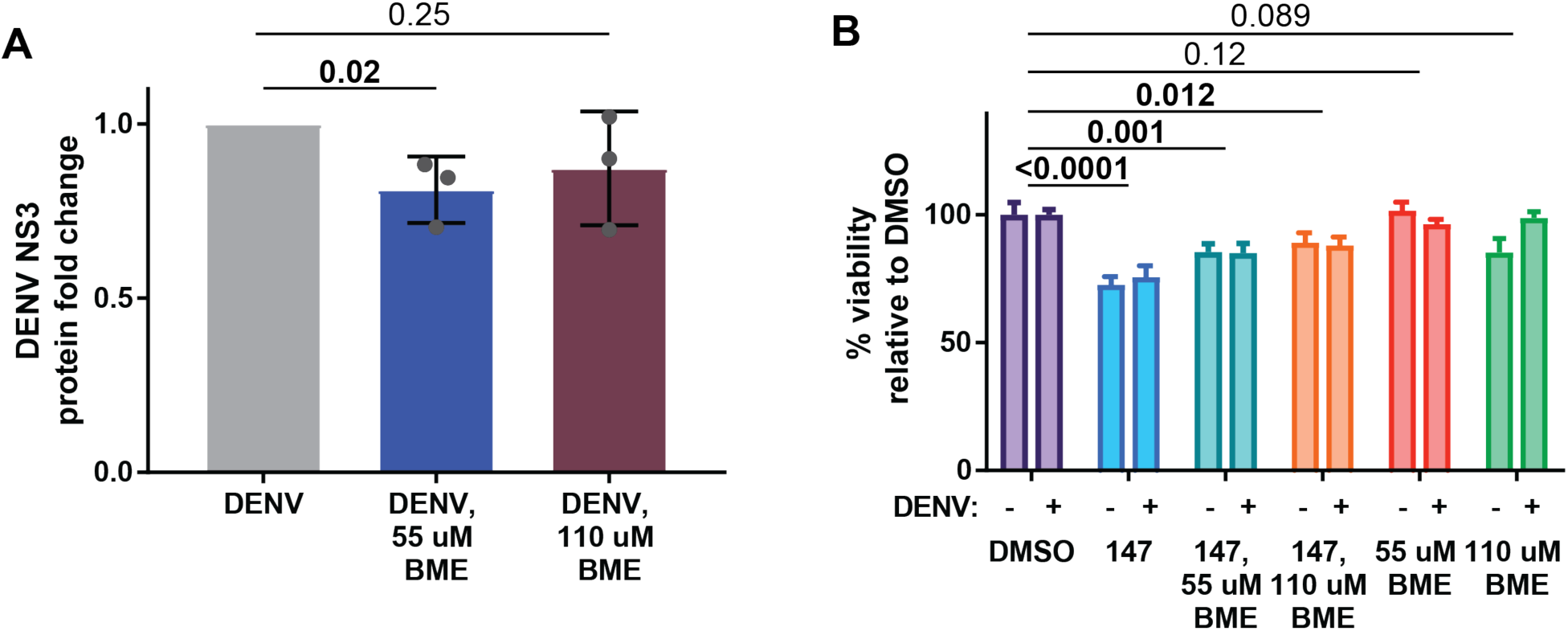
BME addition does not lower DENV infection or impair cell viability. A. Western blot data showing BME does not reduce DENV propagation in Huh7 cells. Cells were pretreated for 16h, then infected (where indicated) with DENV-2 strain BID-V533 at MOI of 3 for 3 hours. Media and treatments were replaced, and cells were collected 24hpi. Cells were lysed, and protein samples were run out on SDS-PAGE gels and visualized using PVDF membranes. B. Cell viability data showing 147/BME treatment only modestly affects Huh7 cell viability over the course of treatment. Cells were pretreated for 16h, then infected (where indicated) with DENV-2 strain BID-V533 at MOI of 3 for 3 hours. Media and treatments were replaced, and viability was measured 24hpi using the Promega Cell Titer Glo reagent. Graphs represent three to six biological replicates. Error bars represent SEM.

**Figure 5 – supplement 1.**
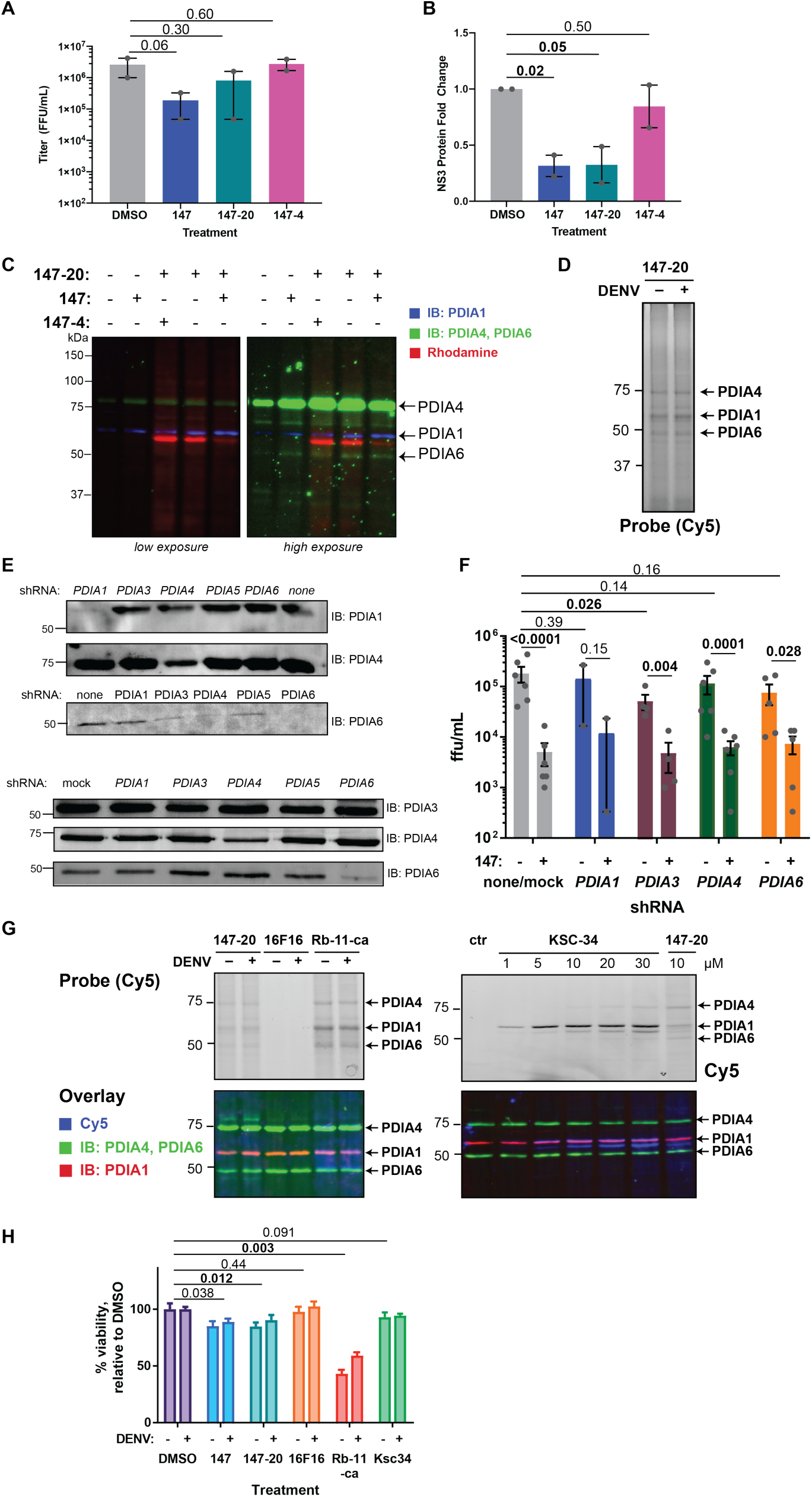
PDIs are covalent targets of 147 but knockdown or inhibition of PDIs are does not recapitulate the reduction in DENV viral infection. A. Titer data showing 147-20 has a similar effect on virus replication as the parent compound 147. In contrast, compound 147-4 has no observable effect on viral titer levels. Cells were pretreated with 10 µM of the respective compound for 16h, then infected with DENV-2 strain BID-V533 at MOI of 3 for 3 hours. Media and treatments were replaced and titer levels were measured 24 hpi. B. Western blot quantification of DENV NS3 levels after treatment with 147, 147-20, or 147-4. Cells were pretreated with 10 μM respective compound for 16h, then infected with DENV-2 strain BID-V533 at a MOI of 3 for 3 hours. Media and treatments were replaced, and cells were collected at indicated timepoints. Cells were lysed, and protein samples were run on SDS-PAGE gels and visualized using PVDF membranes. C. Western blot showing overlay of TAMRA and probes for PDIs. Huh7 cells were treated with indicated compounds at 3 μM (for single compound treatments) or 3 μM 147-20 and 9 μM 147/147-4 (for double compound treatments) for 16-18 hours before harvest. Cells were lysed, protein concentration normalized, and Click chemistry was performed using a TAMRA-biotin-azide trifunctional probe for 1 hour at 37°C. Samples were run on SDS-PAGE gels and targets were visualized on PVDF membranes. D. Representative western blot showing the addition of DENV does not result in additional target labeling of **147-20**. Huh7 cells were treated with 10 μM **147-20** for 16-18 hours before harvest, then infected with DENV2 strain BID-V533 at MOI of 3 for 3 hours. 24 hpi, cells were lysed, normalized to 1 mg/mL protein, and Click chemistry was performed using a Cy5 azide fluorophore for 1 hour at 37°C. Samples were immediately run on SDS-PAGE gels and targets were visualized on PVDF membranes. E. Western blot showing successful creation of PDI knockdown cell lines. Lentiviral vectors were produced using a pooled combination of 2 or 3 shRNAs in HEK293T cells. Lentiviruses were transduced into Huh7 cells, and knockdown cells were selected using puromycin. Polyclonal knockdown cells were maintained as normal Huh7 cells. Two independent batches of knockdown cell lines were produced; the top three panels represent the first, the bottom four panels represent the second. F. PDI knockdowns cells exhibit a similar response to **147** as the parental Huh7 cell line or mock-transduced cells. Only PDIA3 knockdown results in a small but significant reduction. Cells were treated with **147** (where indicated) for 16 hours, then infected with DENV-2 strain BID-V533 at MOI 3 for 3 hours. Media and treatments were replaced, and media was harvested 24hpi. Titer assay was performed as described. Graphs represent 2-6 independent biological replicates. Error bars represent SEM and p-values from ratio paired t-tests are shown. G. Western blot showing overlay of Cy5 fluorophore (from Click chemistry) and antibody probes for PDIs. Huh7 cells were treated with indicated compounds (10 μM **147-20**, 10 μM **16F16**, 20 μM **Rb-11-ca**, or 5 μM **KSC34)** for 16-18 hours before harvest. Cells were lysed, normalized to 1 mg/mL protein, and Click chemistry was performed using a Cy5 azide fluorophore for 1 hour at 37°C. Samples were immediately run on SDS-PAGE gels and targets were visualized using PVDF membranes. **16F16** lacks a click handle preventing fluorophore labeling. H. Cell viability data with PDI inhibitors. **147** only minimally reduced viability. **16F16** and **KSC34** do not impact viability, but **Rb-11-ca** leads to a greater than 50 % reduction in viability. Cells were pretreated with 10 μM **147**, 10 μM **16F16**, 20 μM **Rb-11-ca**, or 5 μM **KSC34** for 16h, then infected (where indicated) with DENV-2 strain BID-V533 at MOI of 3 for 3 hours. Media and treatments were replaced, and viability was measured 24hpi using the CellTiter-Glo reagent. Graphs represent three biological replicates. Error bars represent SEM.

**Figure 6 – supplement 1.**
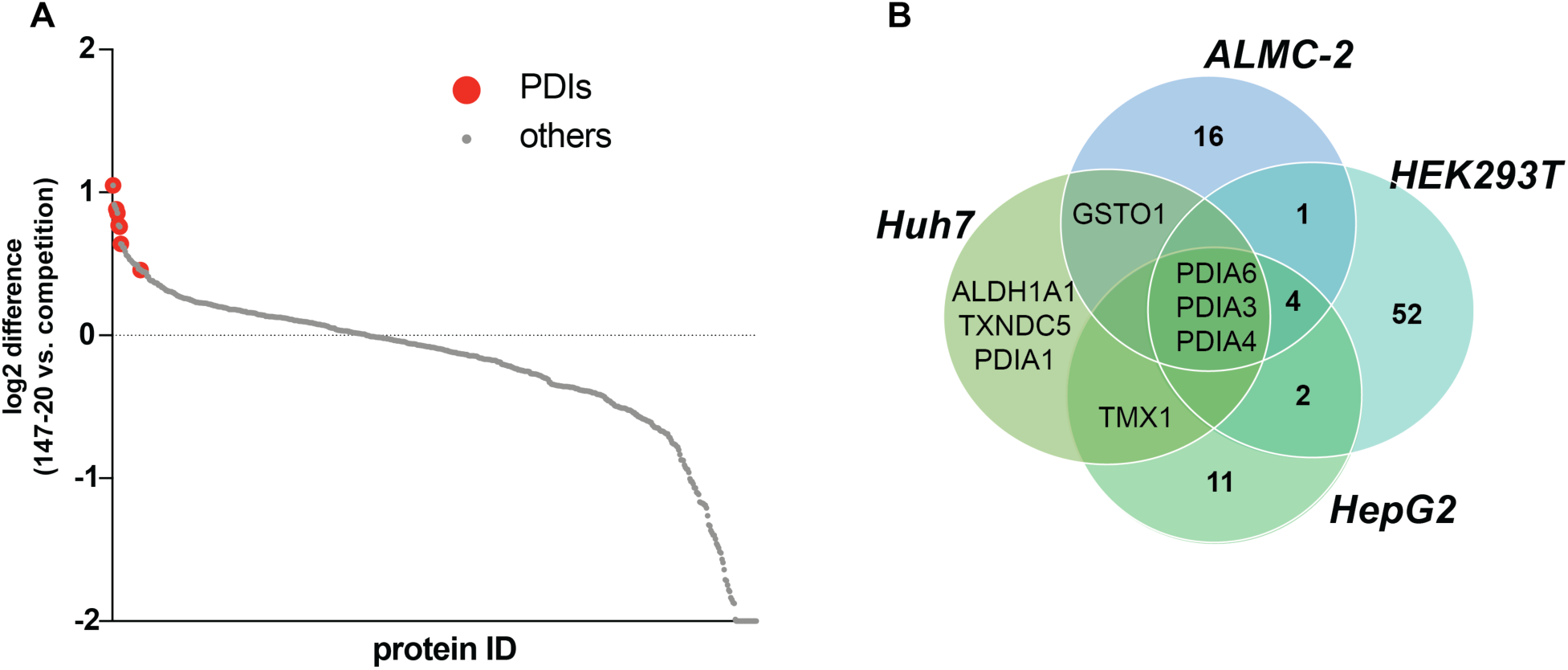
Identification of covalent protein target of 147 in Huh7 cells. A. Chemoproteomic target identification for **147-20** shows an enrichment of several PDIs compared to the **147**-competition control. Quantitative proteomics using TMT-11plex reagents was used to compare log2 intensities across 4 replicates of probe (**147-20** alone) versus probe in competition with excess parent compound **147**. Targets were ordered from most enriched (high log2 fold change) to least enriched (low log2 fold change). Protein disulfide isomerases (PDIs, highlighted as red dots) were among the most enriched targets identified. B. Venn diagram of highly enriched targets found in Huh7 cells in this study compared to HEK293T, HepG2, and ALMC-2 cells in a previous study (Paxman et al., 2018). Previous target ID studies were comparable to the study conducted herein.

**Figure 7 – supplement 1.**
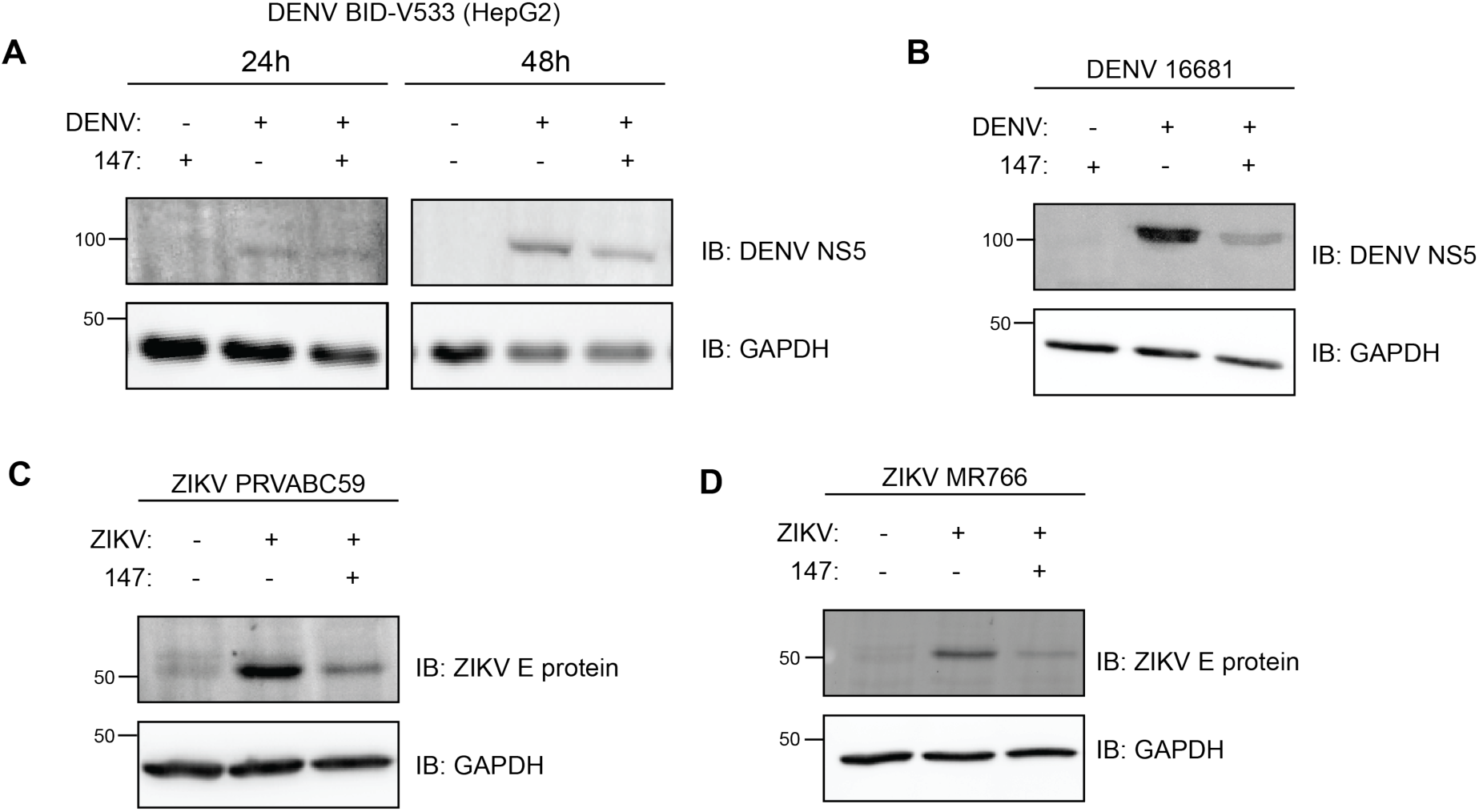
Reduction of viral proteins in DENV and ZIKV strains by compound 147. A. Representative western blot showing similar phenotype in HepG2 cells as in Huh7 cells. HepG2 cells were pretreated with 10 μM **147** for 16h, then infected with DENV-2 strain BID-V533 at MOI of 3 for 3 hours. Media and treatments were replaced, and cells were collected at 24 and 48 hpi. Cells were lysed, and protein samples were run on SDS-PAGE gels and visualized using PVDF membranes. GAPDH is included as a loading control. B. Representative western blot showing attenuation of DENV-2 16681 viral protein levels on treatment with 147. Cells were pretreated with 10 μM 147 for 16h, then infected with DENV-2 strain 16681 at MOI of 3 for 3 hours. Media and treatments were replaced, and cells were collected at 24hpi. Cells were lysed, and protein samples were run on SDS-PAGE gels and visualized using PVDF membranes. GAPDH is included as a loading control. C. Representative Western Blot showing attenuation of ZIKV PRVABC59 viral envelope (E) protein levels on treatment with 147. Cells were pretreated with 10 μM 147 for 16h, then infected with ZIKV strain PRVABC59 at MOI of 0.5 for 1 hour. Media and treatments were replaced, and cells were collected at 24hpi. Cells were lysed, and protein samples were run on SDS-PAGE gels and visualized using PVDF membranes. GAPDH is included as a loading control. D. Representative Western Blot showing attenuation of ZIKV MR766 viral E protein levels on treatment with 147. Cells were pretreated with 10 μM 147 for 16h, then infected with ZIKV strain MR766 at MOI of 0.5 for 1 hour. Media and treatments were replaced, and cells were collected at 24hpi. Cells were lysed, and protein samples were run on SDS-PAGE gels and visualized using PVDF membranes. GAPDH is included as a loading control.

